# Neural Stem Cell Relay from B1 to B2 cells in the adult mouse Ventricular-Subventricular Zone

**DOI:** 10.1101/2024.06.28.600695

**Authors:** Arantxa Cebrian-Silla, Marcos Assis Nascimento, Walter Mancia, Susana Gonzalez-Granero, Ricardo Romero-Rodriguez, Kirsten Obernier, David M Steffen, Daniel. A. Lim, Jose Manuel Garcia-Verdugo, Arturo Alvarez-Buylla

## Abstract

Neurogenesis and gliogenesis continue in the Ventricular-Subventricular Zone (V-SVZ) of the adult rodent brain. B1 cells are astroglial cells derived from radial glia that function as primary progenitors or neural stem cells (NSCs) in the V-SVZ. B1 cells, which have a small apical contact with the ventricle, decline in numbers during early postnatal life, yet neurogenesis continues into adulthood. Here we found that a second population of V-SVZ astroglial cells (B2 cells), that do not contact the ventricle, function as NSCs in the adult brain. B2 cell numbers increase postnatally, remain constant in 12-month-old mice and decrease by 18 months. Transcriptomic analysis of ventricular-contacting and non-contacting B cells revealed key molecular differences to distinguish B1 from B2 cells. Transplantation and lineage tracing of B2 cells demonstrate their function as primary progenitors for adult neurogenesis. This study reveals how NSC function is relayed from B1 to B2 progenitors to maintain adult neurogenesis.

## Introduction

Stem cells, primary progenitors with the capacity to self-renew and generate multiple cell types, have been shown to persist postnatally in many tissues. These cells play a key role in tissue homeostasis and regeneration. Although the brain was once considered an exception, it is now well recognized that a population of cells with astroglial properties derived from radial glia (RG) can function as neural stem cells (NSCs) in the adult brain^1–3^. These cells are enriched in the Ventricular-Subventricular Zone (V-SVZ), an extensive germinal niche in the walls of the lateral ventricles^4,5^. NSCs in the rodent V-SVZ serve as primary progenitors for the generation of large numbers of neurons destined for the olfactory bulb^6,7^ and a smaller number of oligodendrocytes that migrate primarily into the corpus callosum^8–11^. The V-SVZ NSCs have been identified as a subpopulation of astrocytes (also known as B cells) derived from RG during early development^12–15^. Like ventricular RG, B1 cells retain neuroepithelial properties, with an apical domain that has a primary cilium projecting into the ventricle^4,13,15,16^. However, unlike RG, B1 cells’ apical contacts are surrounded by ependymal cells forming pinwheels^4^. Interactions with this epithelium and factors derived from the cerebrospinal fluid (CSF) have been suggested to regulate neurogenesis and B1 cells’ NSC function^17,18^.

Whereas a small subpopulation of B1 cells self-renew, the majority divide symmetrically to generate young neurons, which results in a rapid decrease in B1 cell number with age^19,20^. While neurogenesis also decreases with age, the rate at which olfactory bulb neuron production decreases is significantly lower than the rate at which B1 cells are depleted ^19^. This suggests that a second population of progenitors contributes to olfactory bulb neurogenesis into adulthood. An initial characterization of the V-SVZ identified a second population of B cells (B2 cells) that share marker expression and ultrastructural features with B1 cells but are generally located more basally in the V-SVZ^21^. Like B1 cells, B2 cells have ultrastructural features of astroglia and express proteins typically present in astrocytes, like glial fibrillary acidic protein (GFAP), Vimentin, and Nestin^22^. Unbiased single-cell RNA-sequencing studies of the V-SVZ have not revealed differences between B1 and B2 cells^23–32^ suggesting that these cells share many molecular properties. Interestingly, some B2 cells incorporate ^3^[H]-thymidine, suggesting they proliferate and possibly serve as progenitor cells^21^. A recent study also suggests that basal progenitors can function as NSCs in the V-SVZ^33^. However, whether B2 cells can function as neuronal and glial progenitors or whether they correspond to a population of more differentiated parenchymal astrocytes, remains controversial^33,34^.

Here, we show that B2 cells serve as a reservoir of primary progenitors in the V-SVZ, which explains how neurogenesis is maintained as B1 cells are depleted in adult mice. Together, our results indicate that NSCs function is progressively relayed from B1 to B2 cells in the adult mammalian brain.

## Results

### B2 cells are non-apical V-SVZ astrocytes

The identification of B2 cells is challenging given that these cells have similar morphology, ultrastructure, and marker expression to B1 cells. B1 cells can be identified by their thin apical extension with a primary cilium that contacts the ventricle^4^. Although B2 cells are found beneath the ependymal layer, B2 cells might contact the ventricle through thin processes. To visualize thin processes of B2 cells, we used serial section reconstructions and transmission electron microscopy (TEM). Full reconstructions of 41 B2 cells at P10, 17 at P60, and 11 at P365 showed that B2 cells did not have an apical contact that reached the ventricle. However, in contrast to a recent study^33^, all these 69 serially reconstructed B2 cells had primary cilium. The primary cilium and basal body of B2 cells were always located away from the ventricular surface (Figures 1. A-E and S1A-J). B2 cell’s primary cilium had 9+0 microtubule organization and showed ciliary pockets and occasionally ciliary rootlets (Figure S1A, B). B1 and B2 cells showed similar cytoplasmic and nuclear organization^35^, contacted blood vessels and fractones^36–38^, and ensheathed intermediate progenitors (C cells) and neuroblasts (A cells)^6,21^ (Figures 1A, B and S1A-G). Many B2 cells were found very close to ependymal cells (e.g. Figure 1A, B2 cells were < 5µm from the ventricular surface) and cannot be only identified by their location as recently suggested^33^. In summary, the germinal layer on the walls of the mouse lateral ventricles has 2 domains: an apical ventricular zone (VZ) bordering the ventricle and a non-apical subventricular-zone (SVZ) domain deeper in the tissue between the VZ and the striatum ^39^. Therefore, the location of the primary cilium and basal body provides an objective way to distinguish B2 from B1 cells in the V-SVZ.

**Figure 1.**
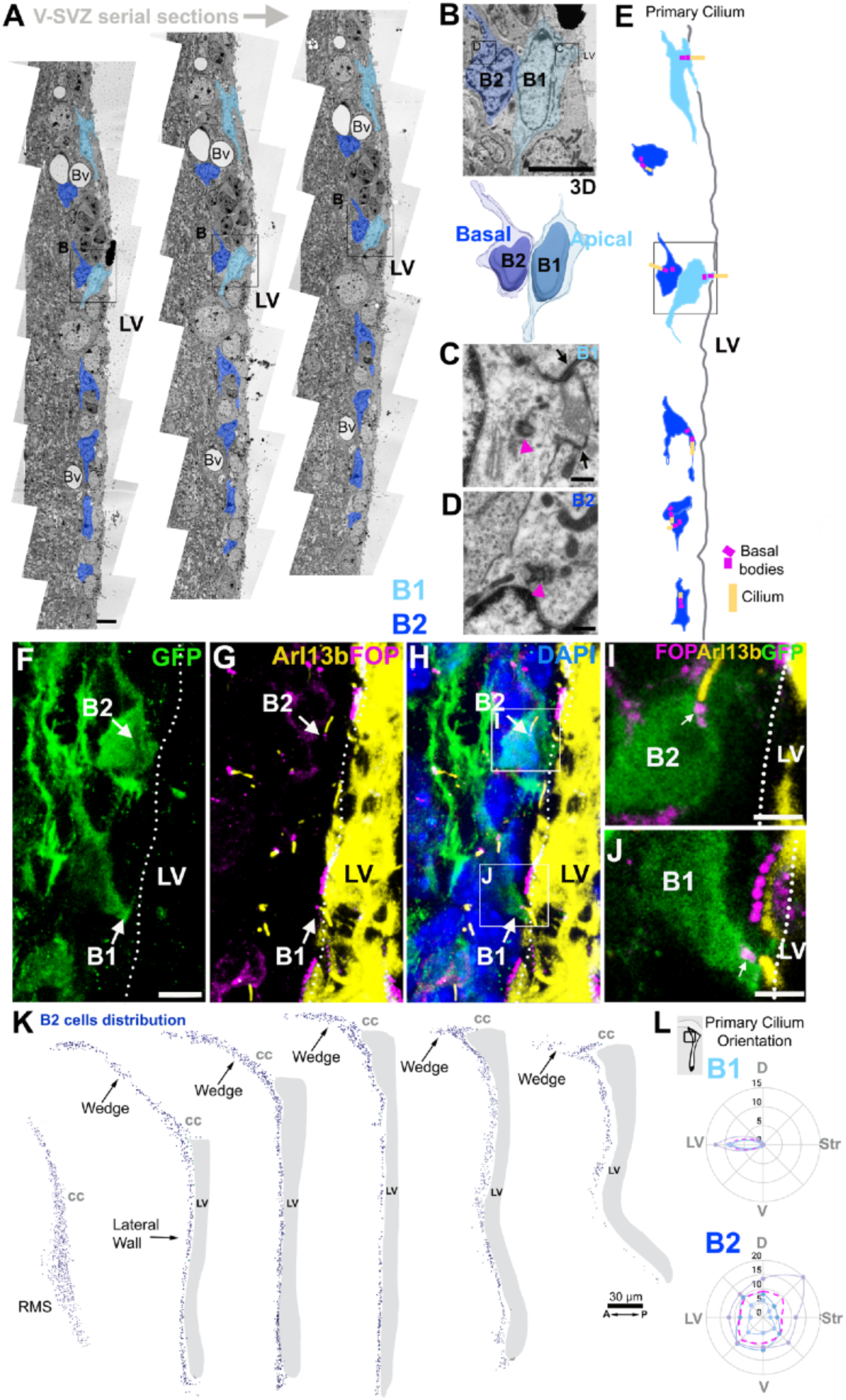
B2 cells are non-apical V-SVZ astrocytes. (**A**) Transmission electron microscopy (TEM) serial sections of the P60 V-SVZ (3 ultrathin sections spaced 1.7μm (3 out of 150 studied)). B1 and B2 cells were pseudo-colored in light and dark blue, respectively. B1 cells contact the lateral ventricle through a small apical contact. Note that B2 cells lack contact with the lateral ventricle and can be located adjacent to ependymal cells. (**B**) High magnification of B1 and B2 cells in (A) and their three-dimensional reconstruction. (**C, D**) High magnification micrographs of B1 and B2 cells’ basal bodies. (**C**) B1 cell apical contact, identified by the presence of tight junctions (arrows) with ependymal cells, shows a basal body (arrowhead). (**D**) B2 cells lack tight junctions, and the basal body (arrowhead) is basally located. See full B1 and B2 cells primary cilium-basal body reconstruction (Figure S1H-J). (**E**) Summary schematic of the V-SVZ TEM serial reconstruction showing the localization of the primary cilium (yellow) and basal bodies (magenta) in B1 (light blue) and B2 (dark blue) cells. (**F-H**) Confocal images of a V-SVZ coronal section showing GFP^+^ B1 and B2 cells and their primary cilium-basal bodies labeled with Arl13B (yellow) and FOP (magenta). Dotted lines outline the apical surface. (**I-J**) High magnification images from (H) showing a GFP^+^ B2 cell with a primary cilium-basal body basally located and lacking an apical contact. The B1 cell shows its primary cilium-basal body contacting the ventricle. (**K**) Map of the distribution of B2 cells in the V-SVZ through anterior to posterior levels. Note that B2 cells are also present in the wedge, a region devoid of ventricular cavity. (**L**) Radar maps of dorsal B1 and B2 cells’ primary cilium orientation. Average of B2 cells primary cilium localization (magenta) shows random orientation (n=3, 427 cells). See also Figures S1 and S2. Bv: Blood vessel, CC: corpus callosum, RMS: Rostral Migratory stream, LV: Lateral Ventricle. Scale bars: 5μm (A, F-H), 2μm (B), 250nm (C-D), 2.5μm (I-J) and 30 μm (K).

Determining the overall distribution and number of B2 cells would not be practical using TEM. We therefore developed a strategy to detect the primary cilium-basal body among B cells using serial z-plane confocal microscopy in hGFAP::GFP P60 mice. GFP^+^ B cells ^40^ with basally located primary cilium were identified as B2 cells. The primary cilium was identified by Arl13b staining and the basal body stained with Ɣ-tubulin antibodies (Figures 1 F-H and S1K). FGFR1 Oncogene Partner (FOP) expression in B cells confirmed the location of the primary cilium-basal body (Figure 1F-J). To distinguish B2 cells from parenchymal astrocytes, we used S100a6 staining; S100a6 is highly expressed in B1 and B2 cells, but not in striatal astrocytes (Figure S2A-H) ^41^. B2 cells were identified as S100a6^+^/GFP^+^ with their primary cilium in the SVZ and away from the ventricular surface (Figure S2B). While B1 cells’ primary cilium was at the apical surface and oriented towards the ventricle, B2 cells’ primary cilium was oriented in multiple directions, deeper in the tissue (Figures 1L and S1L). In serial coronal sections of the V-SVZ, B2 cells were present at all rostro-caudal levels. B2 cells’ soma could be observed as close as 5µm from the ventricular surface (right beneath ependymal cells) and as far as 0.5 mm in the dorsolateral extension wedge region (Figures 1K and S2G-J). Thus, serial z-plane confocal microscopy allows for the identification of B2 cells and shows that the wedge, which does not contain ventricular surface and B1 cells, is rich in B2 cells.

### The number of B2 cells increases postnatally as B1 cells decrease

In whole-mount preparations^4,16^, and using serial en-face z-plane confocal images from hGFAP::GFP mice, we identified B2 cells based on the basal location of the primary cilium-basal body (Figure 2A-F). ß-catenin staining was used to visualize the apical epithelial contacts. B1 cells’ primary cilia were identified at the ventricular surface in the center of pinwheels; however, B2 cells’ primary cilia were deeper in the tissue beneath the ependymal layer but within the high cell density of the SVZ. We mapped the distribution of B1 and B2 cells in whole-mounts of the lateral wall of the lateral ventricles at P60 (Figure 2G, H). At this age, approximately half of the B cell population in the V-SVZ corresponded to B2 cells (17,246 B2 cells and 15,864 B1 cells). These results show that B2 cells are as common as B1 cells in young adults and are widely distributed in the V-SVZ extending into the wedge.

**Figure 2.**
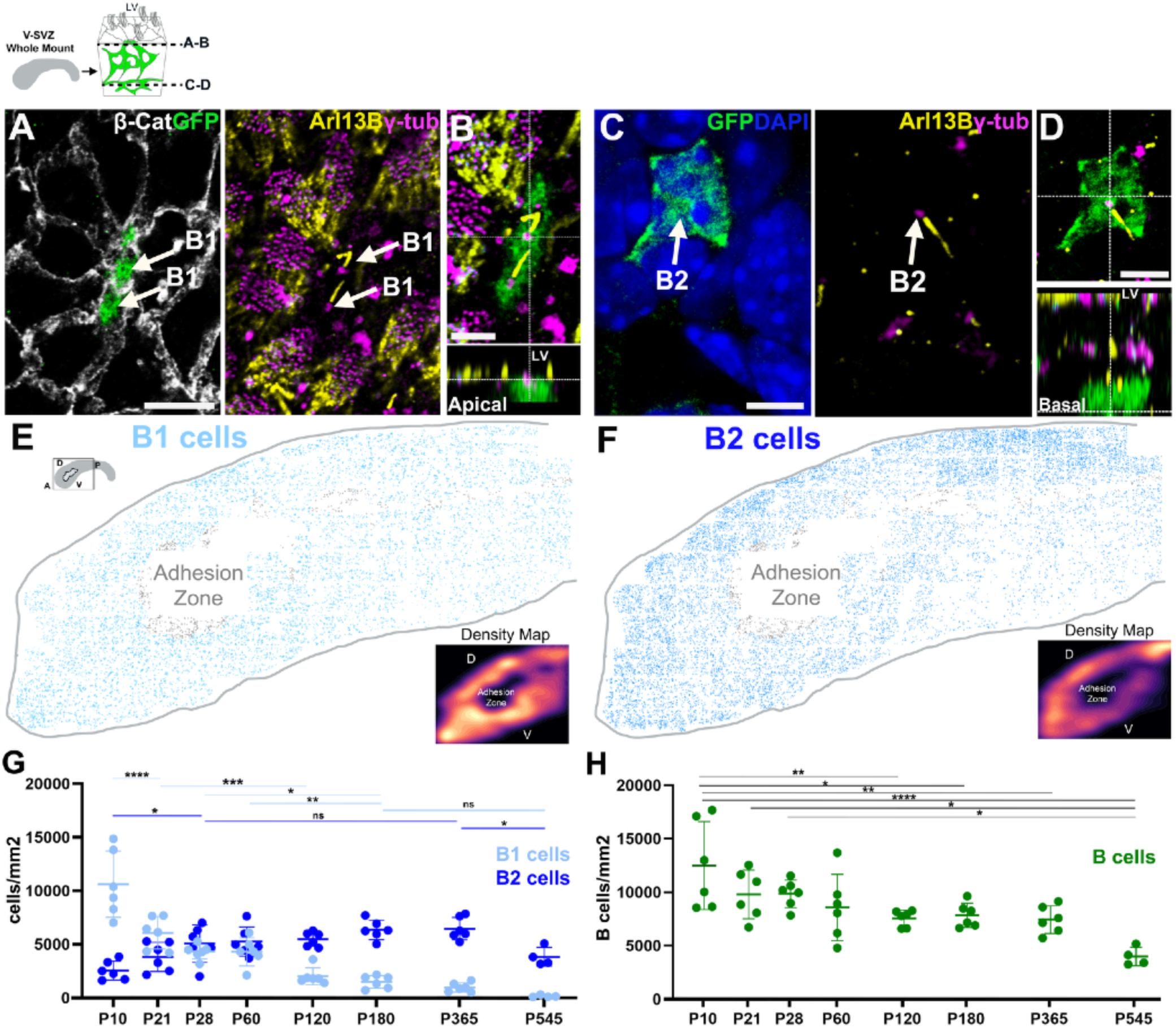
Large-scale identification of B1 and B2 cells. (**A-D**) Confocal images of a V-SVZ whole-mount preparation from hGFAP::GFP mouse. (**A, B**) Images of the surface of the lateral ventricle showing B1 cells apical contact, identified by GFP^+^ expression at the center of the pinwheels and the apical localization of their primary cilium (Arl13b, yellow) and basal body (γ-tubulin, magenta). (**C, D**) Images obtained deeper into the SVZ reveal B2 cells. Cells were identified by GFP expression and the basal location of their primary cilium-basal body. (**E, F**) Maps of B1 (15,864 cells, light blue) and B2 (17,246 cells, dark blue) cells in a P60 female whole-mount preparation. Density Maps show the overall distribution of B1 and B2 cells. (**G**) Quantifications of B1 and B2 cells over time in V-SVZ whole mounts (n=6, 3 females and 3 males/time point, except P545 3 females and 1 male). B1 cells decreased sharply within the first 4 months of age (from 10625 to 2015 GFP^+^ cells/mm2, p-value <0.0001). B2 cell numbers increased between P10 and P28 (from 2547 to 5067 GFP^+^ cells/mm2, p-value 0.0132) and these numbers remained relatively stable up to P365 (6465 GFP^+^ cells/mm2, non-significant). B2 cell numbers decreased significantly from 6465 to 3832 GFP^+^ cells/mm2 between P365 and P574, p-value 0.025. (**H**) Quantifications of B1 and B2 cells (B cells) over time. The pool of B cells decreases with age from 12499 (P10) to 3994 (P545) cells/mm2, p-value <0.0001. See also Figure S3. LV: lateral ventricle. Scale bars: 20μm (A) and 10μm (B-D).

To compare how B1 and B2 cell populations change with age, we analyzed whole-mount preparations from hGFAP::GFP mice at P10, P28, P60, P120, P240, P365 and P547 (n=3/age/ sex/region). B1 and B2 cells were identified by the location of the primary cilium-basal body (see above). Quantifications showed the numbers of B2 cells increased significantly in juvenile mice (P10 to P28, from 2547 to 5067 GFP^+^ cells/mm2, p-value 0.0132), and the population remained constant into adult life (up to P365) (Figure 2G). However, in aged mice (P547) the number of B2 cells decreased to almost ½ of the number observed at one year (from 6465 to 3832 cells/mm2, p-value 0.025) (Figure 2G). In contrast, and consistent with previous observations^19^, the number of B1 cells decreased during the first months of age with very few B1 cells remaining by 1.5 years of age (p-value: 0.0001)(Figure 2G). This pattern was similar in the antero-ventral (AV) and postero-dorsal (PD) regions (Figure S3A-C), both regions with high levels of adult neurogenesis ^4^. We did not observe significant sex differences in the number of B1 or B2 cells at the different ages studied (Figure S3D-F). When combining B1 and B2 cells, we found that the overall number of B cells declined by one-third with age (from 12499 cells/ mm2 at P10 to 3994 cells/mm2 at P545) (Figure 2H). These results show that B2 cells become the predominant B cell population after 2 months of age.

### Large numbers of B2 cells are postnatally derived from B1 cells

The number of B2 cells increases significantly between P10 and P28. To determine if in these juvenile animals B2 cells are generated from B1 cells, we labeled B1 cells by a single injection of Violet Cell Trace (VCT) into the lateral ventricles of P14 hGFAP:GFP mice (Figure 3A). This tracer is an ester that is intracellularly converted to a fluorescent derivative that binds to amine groups in proteins, resulting in long-term dye retention ^42^. Cells in contact with the CSF were labeled by the VCT injection. To avoid any intra-parenchymal labeling from the injection track and ensure that only cells exposed to the CSF became labeled, we only analyzed the contralateral side to the injection side. The wedge region, which does not contact the lateral ventricle, was not labeled at any of the studied timepoints (1, 7 and 30 days after injection (n=3/time point)). At these time points, we found strong VCT labeling in apical cells (ependymal and B1 cells) and no evidence of diffusion of dye into the SVZ (Figure 3B).

**Figure 3.**
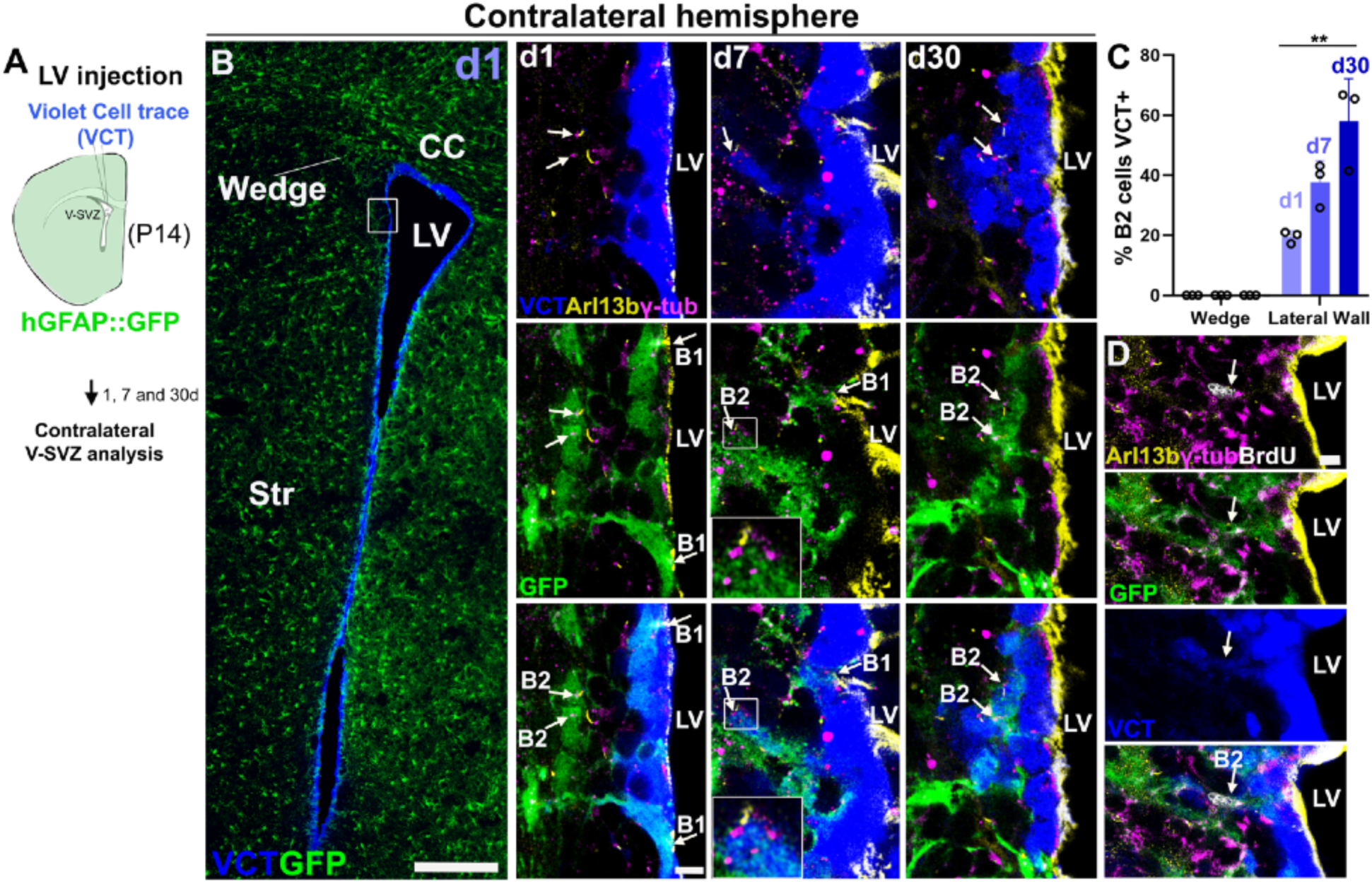
B1 cells give rise to B2 cells. (**A**) Schematic of Violet Cell Trace (VCT) injection into the lateral ventricles in P14 hGFAP:GFP mice. (**B, C**) Confocal images of the contralateral lateral ventricle 1 day after injection, showing VCT is restricted to the V-SVZ and does not diffuse freely into the brain parenchyma. B2 cells labeled with VCT, identified by GFP^+^ expression and the basal localization of their primary cilium (Arl13b, yellow) and basal body (γ-tubulin, magenta), significantly increased at d30 (from 19.45% to 57.93%, p-value 0.0056). No labeled cells were found in the wedge extension. (**D**) Confocal images of a B2 cell positive for VCT and BrdU. See also Figure S3. CC: corpus callosum, LV: lateral ventricles. Scale bars: 250μm (B) and 5μm (B, D).

Using the criteria of primary cilium-basal bodies location, we quantified the number of B2 cells labeled with VCT over time after injection. We found that the proportion of VCT^+^ B2 cells significantly increased from d1 to d30 after injection (from 19.45% ± 2.00 to 57.93 ± 14.24, p-value: 0.005). This suggests that B1 cells give rise to B2 cells in the lateral wall of juvenile mice. To further investigate if this conversion was associated with cell division, we injected BrdU immediately after intraventricular VCT administration and analyzed the contralateral lateral wall 30 days later (n=4) (Figure S3G). BrdU-labeled B1 or B2 cells were identified based on the location of primary cilium-basal bodies in coronal sections (8-10) of the V-SVZ from hGFAP:GFP mice (n=4). Of the 284 BrdU^+^ cells analyzed, 18 were B1 cells and 181 were B2 cells (Figure S3H). Interestingly, almost one-third (32.48% ± 9.09) of the BrdU-labeled B2 cells were also labeled by VCT (Figures 3C and S3I, J). These findings suggest that in juvenile animals, B1 cells give rise to B2 cells, and that this conversion may be linked to cell division.

### Transcriptomic profiling of apical (B1) and non-apical (B2) cell identities

Given that B1, but not B2 cells, are part of the VZ epithelium, we hypothesized that B1 cells must differ from B2 cells at the transcriptomic level. Analyses of published single-cell transcriptomic datasets of the V-SVZ identified the activation state^24,25,27–32^ and regionality ^23,24,26,43,44^ as the main sources of transcriptomic variance among B cells. Previous studies have suggested that VCAM1 and Prominin1 are primarily expressed by B1 cells ^33,45^. However, analysis of *Vcam1 and Prom1* expression in our V-SVZ single-cell RNA sequencing (scRNA-seq) analysis ^23^ showed that *Vcam1* was expressed widely in B cells (Figure S4A,B). *Prom1* expression was also found throughout the B cell clusters and was also present in C and A cells. Therefore, neither *Vcam1* nor *Prom1* mRNA levels were specific to B1 identity (Figure S4A, B). RNAscope for *Vcam1* or *Prom1* in hGFAP::GFP mice, together with localization of the primary cilia-basal body, confirmed that *Vcam1* and *Prom1* mRNAs were present in both B1 and B2 cells, including cells in the wedge. *Prom1* was also expressed in C and A cells (Figure S4C-F).

To identify B1 cells for scRNA-seq analysis of the V-SVZ, we genetically tagged cells contacting the ventricle before dissociation and single-cell transcriptomic analysis. An adenovirus encoding for GFP and Cre (Ad-GFP-Cre) was injected into the lateral ventricles of mice carrying a floxed Tdtomato gene (Figure 4A). Twenty-four hrs after intraventricular injection, whole mounts were prepared from the contralateral hemisphere for analysis. We found that 23.83% ± 6.89 of B1 cells and 61.64% ± 23.14 ependymal cells were TdTomato^+^; B2 cells identified by the basal location of their primary cilium-basal body were Tdtomato^-^ (n=3, Figure 4B). Consistent with the absence of B1 cells in this region, the wedge was devoid of Tdtomato labeling (n=3, Figure 4C).

**Figure 4.**
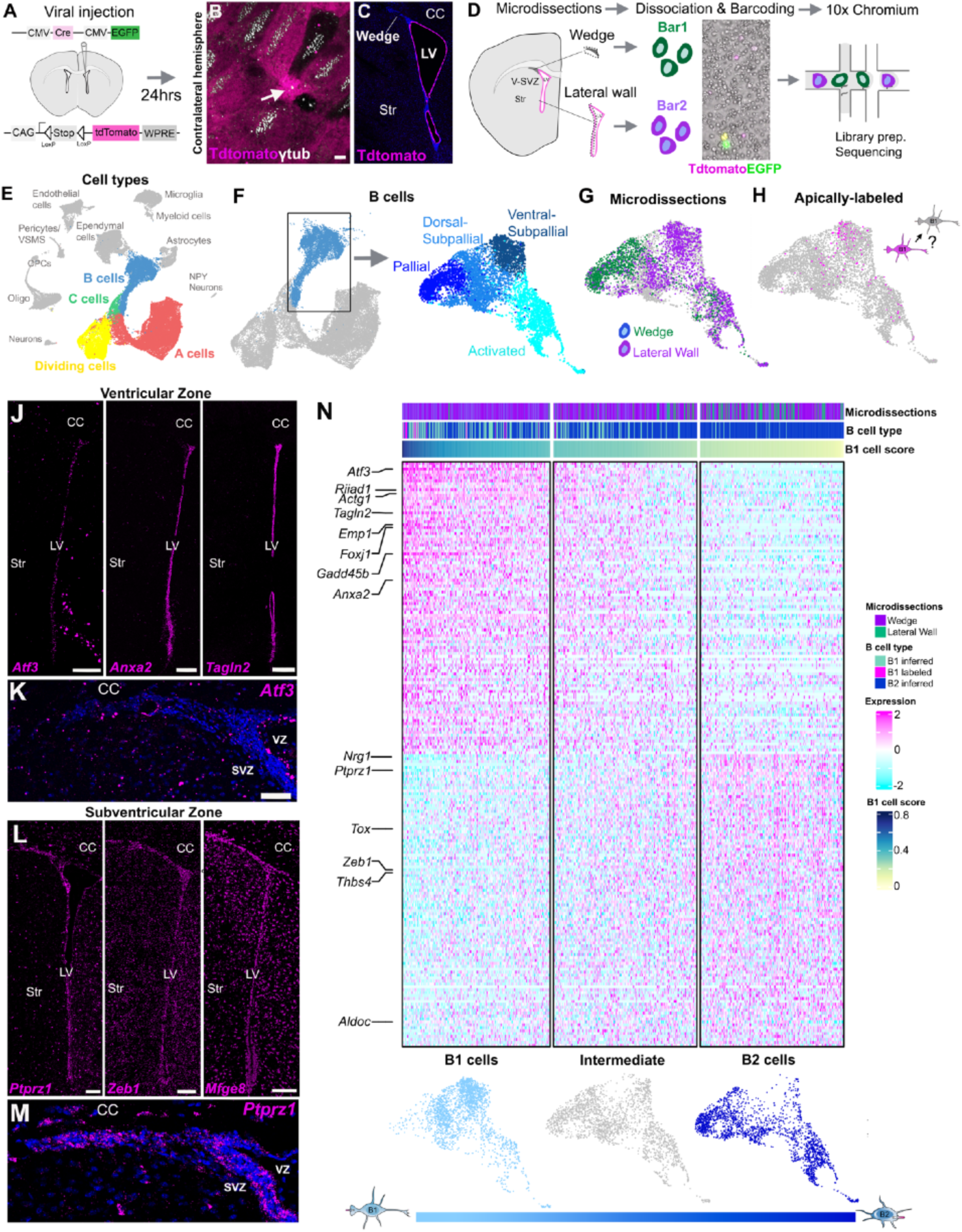
B1 and B2 cell transcriptomic profiles. (**A**) Schematic of the intraventricular injection of Adenovirus;Cre, GFP into Tdtomato mice. (**B-C**) Confocal images of the contralateral V-SVZ 24 hrs after injection. Note that a subpopulation of B1 cells and ependymal cells, identified by their basal bodies (γ-tubulin, white), were Tdtomato^+^. The wedge region did not show Tdtomato labeled cells. (**D**) Schematic of the whole-cell single-cell isolation and sequencing protocol (scRNA-Seq). The wedge region and lateral wall were microdissected from the contralateral hemisphere 24 hrs after injection. Cells were dissociated, incubated with MULTI-seq barcodes, multiplexed, and loaded into the 10x Chromium Controller. (**E**) UMAP plot of scRNA-Seq cell types captured after sequencing and downstream analysis. (**F**) UMAP plot showing subset and re-clustered B cells. B cell clusters annotated by their activation states and regional identities. See also Figure S4. (**G**) Distribution of barcoded cells from the Lateral wall and wedge microdissections in B cell clusters. (**J**) Distribution of apical-labeled cells (B1 cells) identified by the expression of *Tdtomato, Cre* and *GFP.* (**J-K**) Confocal images depict RNA expression of B1 cells differentially expressed genes *Atf3*, *Anxa2* and *Tagln2* in the ventricular zone (VZ). (**L**) Confocal images depict RNA expression of B2 cells differentially expressed genes *Ptprz1, Zeb1* and *Mfge8* in the subventricular zone (SVZ). **(N)** HeatMap of the differentially expressed genes between B1 and B2 cells. These genes were expressed in a gradient that revealed the prototypical B1 and B2 cell transcriptomic profiles, and identified an intermediate population. See Figures S4 and S5. CC: corpus callosum, Str: Striatum, LV: lateral ventricle. Scale bars: 5μm (B), 100μm (J, L) 30μm (K, M)

For scRNA-seq analysis, we microdissected the wedge region and the lateral ventricle wall from the contralateral hemisphere of 26 Ad-GFP-Cre-injected mice (P28-P30). Cells were dissociated, barcoded, pooled, and loaded onto a 10x chip for single-cell transcriptomic analysis (Figure 4D). Adenoviral vectors can upregulate viral response genes ^46^. In order to mitigate the impact of viral response genes on the clusters of B cells, we integrated our newly generated data with our previously published adenovirus-free dataset from the V-SVZ at the same age ^23,29^. Unsupervised clustering of 47,709 captured cells resulted in discrete groups corresponding to the main cell types within the neurogenic lineage (B-C-A and dividing cells) (Figure 4E). In addition, we observed clusters corresponding to ependymal cells, parenchymal astrocytes, microglial, myeloid, endothelial and mural cells, oligo-progenitors, oligodendrocytes, and neurons (Figures 4E, and S4G, H). As expected, B cells clustered closely with parenchymal astrocytes, but these 2 populations could be distinguished by *Thbs4 and S100b* expression (B cells were *Thbs4^+^/ S100b^-^;* parenchymal astrocytes were *Thbs4*^-^/ *S100b*^+^) ^23,29^(Figure S4H).

To investigate the transcriptomic differences between B1 and B2 cells, we selected and re-clustered B cells (Figure 4F). Consistent with previous works, the clustering of B cells was associated with two main sources of heterogeneity: activation state and regionality. Quiescent (*S100a6^+^Egfr^-^*) B cells could be distinguished from activated B cells (*S100a6^+^Egfr^+^*) ^28,29,47^ (Figure S4I), and subdivided into pallial (*Tfap2c^+^, Crym^-^*), dorsal-subpallial (*Urah^+^, Tfap2c^-^, Crym^-^*) and ventral-subpallial sub-populations(*Urah^-^, Tfap2c^-^, Crym^+^*) ^23,24,44^ (Figures 3F and S4I-K). Cells derived from the wedge and lateral walls had transcriptional profiles consistent with dorsal and ventral identities previously described (Figure 4G).

We identified apically labeled B1 cells by the expression of *Tdtomato*, *Cre,* or *GFP* (Figures 4H and S4K). Since the Ad-GFP-Cre virus only labels ∼25% of B1 cells in the lateral wall, we identified uninfected cells that had the most similar transcriptional profile to infected cells by analyzing their nearest neighbors using Seurat. We inferred that these cells corresponded to unlabeled B1 cells (Figure S4L). The remaining B cells were classified as inferred B2 cells (Figure S4L). To determine if these cells correspond to B1 and B2 cells in the tissue, we conducted a differential gene expression analysis between inferred B1 and B2 cells (Table1, Figure S4M). We selected a set of non-region-specific genes enriched in inferred B1 or B2 cells for RNAscope validation. Inferred B1 cells genes *Atf3, Riiad1, Foxj1, Gadd45b, Emp1,* and *Tagln2* were expressed in B1 and ependymal cells in the VZ, and had low or no expression in B2 cells in the SVZ and the wedge (Figures 4J, K and S5A). An exception was *Emp1,* which is upregulated during mitosis ^48^, was present in dividing cells in the SVZ, including the wedge (Figure S5A, B). We confirmed that *Atf3, Riiad1, Foxj1, Gadd45b, Emp1,* and *Tagln2* were expressed in B1 cells by using RNAscope in *en face* sections of the ventricular wall and co-stained with ß-catenin and Ɣ-tubulin to identify B1 cells’ apical contact.Quantifications showed that 186/320 B1 cells were *Atf3^+^,* 46/80 were *Riiad1^+^,* 91/142 *FoxJ1^+^*, 40/165 *Gadd45b^+^*, 62/180 *Emp1*^+^ and 29/32 *Tagln2*^+^ (Figure S5C, n=2). We also analyzed the expression of non-regional genes differentially expressed in inferred B2 cells. We performed RNAscope for *Ptprz1, Tox, Aldoc, Zeb1, and Mgef8* in combination with Ɣ-tubulin and GFP expression in hGFP:GFP mouse (Figures 4L,M, and S5D). We found that these genes are highly expressed in the V-SVZ, including the wedge region, and confirmed their expression in B2 cells. However, some expression of these genes was also observed in B1 cells (Figure S5D-E).

The histological validation of the 6 genes expressed in B1 cells, and not B2 cells, allowed us to confidently identify B1 cells in our transcriptomic dataset (Figure S5F). By combining the expression of these 6 genes, we scored cells according to the prototypical B1 cell profile (Figure S5G). B cells were found to be on a continuum; while some cells had transcriptomic profiles consistent with prototypical B1 and B2 states, many B cells were found to be in an ambiguous or transitional state (Figure 4N). This analysis reveals sets of genes that identify the transcriptomic profile of prototypical B1 and B2 cells. However, the continuum transition between these two populations, possibly associated with transitional intermediate states, highlights the similarities between these two cell types.

To better understand the functions of proteins encoded by differentially expressed (DE) genes in B1 or B2 cells, we identified possible protein-protein interaction networks among these proteins, and their functional associations, using STRING ^49^. Proteins encoded by DE genes in B1 cells were highly interconnected (4381 interactions among 804 genes), while DE genes in B2 cells were less associated with each other (42 interactions among 303 genes). The top DE genes in B1 cells, *Actg1, Atf3,* and *Anxa2*, represented central nodes that were highly interconnected to several other DE genes. Further analysis reveals that γ-actin (*Actg1*) interacts with Msn, Baiap2, Pfn1, Cfl1, Arpc2, Arpc4, Arpc1b, Rhoa, Actb, Myl12a, Eef1b2, Hspa8. Interestingly, this network is associated with the cytoskeleton and the apical junction complex and is involved in epithelial polarization (Figures 5A and S6A). Atf3 interacts with Junb, Jun, Jund, Fos, Trp53, Ddit3, and Atf4 (Figure S6C), DNA-binding transcription factors, which are part of the AP-1 complex, associated with epithelial function ^50^. Anxa2 also interacted with calcium ion-binding S100 proteins: S100a4, S100a6, S100a10, and S100a11 (Figure S6 D). The Anxa2-S100a10 complex has been associated with the establishment of apical polarity ^51^. We confirmed the expression of Anxa2 in B1 cells’ apical contact using confocal and electron microscopy (Figure S6E-I).

**Figure 5.**
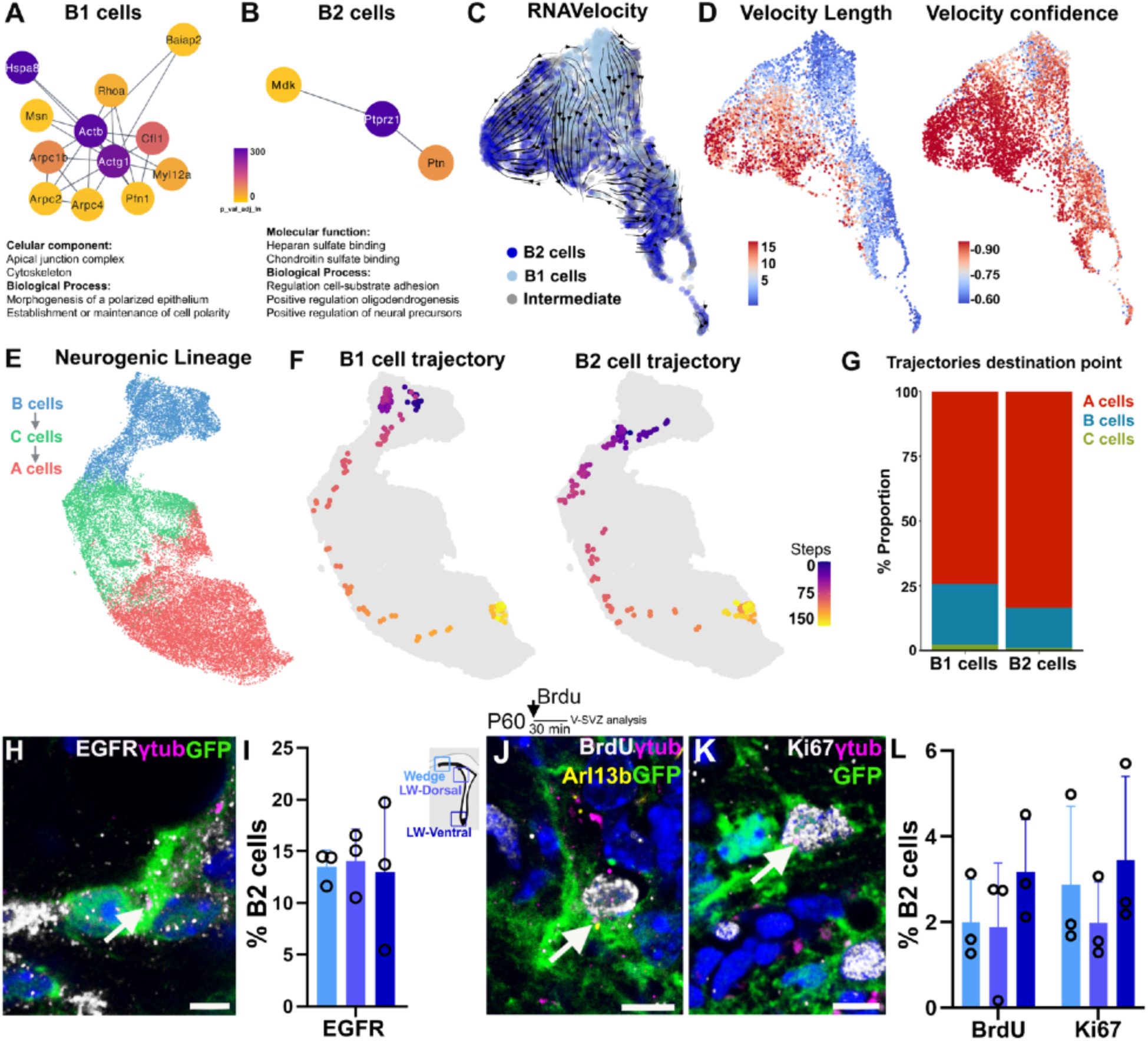
B2 cells are a relay in the neurogenic lineage. (**A-B**) Protein-protein network diagrams of differentially expressed genes in B1 (A) and B2 cells (B). Diagrams show first neighbor interactions for *Actg1* and *Ptprz1* and their functional enrichment annotations. (**C, D**) RNA velocity stream projected onto B cell UMAP plot. Velocity length and confidence for B1 (5.87 and 0.84) and B2 cells (8.32 and 0.91). **(E)** UMAP plot of the neurogenic lineage after cell-cycle regression. (**F-G**) Pseudotime and trajectories of B1 and B2 cells visualized on a 2-dimensional UMAP embedding. Barplots showing the proportion of B1 and B2 cells trajectories ending in B, C or A cells (B1 cells n=2107, B2 cells n= 2049). (**H**) Confocal images of a B2 cell expressing EGFR (white), identified by GFP (green) expression and the basal body location (γ-tubulin, magenta, arrow). (**I**) Percentage of B2 cells EGFR^+^ in the wedge, dorsal, and ventral lateral wall regions (15.32% ± 4.13, 1403 cells, n=3). (**J**) Single dose administration of BrdU and analysis after 30 min in P60 mice. Confocal images of a Brdu^+^ B2 cell (white) in the wedge. (**K**) Confocal images of a Ki67^+^ B2 cell (white) in the wedge. (**L**) Percentage of BrdU+ and Ki67+ B2 cells in the wedge, dorsal and ventral lateral wall regions (BrdU: 2.51% ± 1.43, 5060 cells, n=3; Ki67: 2.92% ± 1.05, 2789 cells, n=3). See also Figure S6. Scale bars: 2.5μm (H) and 5μm (J, K).

These observations are consistent with B1 cells being part of the VZ epithelium and suggest that B1 cells express a set of genes that help maintain their epithelial identity. Among the few protein interactions identified in the differentially expressed genes of B2 cells, one notable network involved the top gene *Ptprz1*, which networks with Ptn and Mdk (Figures 5B and S6B). This network was associated with extracellular matrix binding and was enriched in neural precursor and oligodendrogenesis regulation terms. Interestingly, *Ptn* and *Ptprz1* are markers of outer radial glia, a non-apical neural progenitor found in the developing human brain ^52^.

### B2 cells are relay progenitors in the neurogenic lineage

To better understand B1 and B2 cells activation and transitions, we used RNA velocity ^53^. This method is based on the analysis of mature (spliced) and newly transcribed (unspliced) transcripts to infer the future transcriptomic states of individual cells. We calculated the velocities (i.e. the rate of transcriptomic changes) for B cells. B2 cells had on average a higher velocity compared to B1 cells (8.32 vs 5.87), suggesting B2 cells are more likely to be transitioning to another state compared to B1 cells (Figure 5C, D). Then we inferred the potential B1 and B2 cell lineage trajectories by analyzing their most likely B1 transitions. To mitigate the effects of the cell cycle on cell identity, we re-analyzed the data by regressing out cell-cycle-related genes ^54^. As expected, B1 cells showed a directional flow towards A cells (Figures 5F, G, and S6J). All inferred trajectories starting from a B1 cell transition through a B2 cell state before transitioning into C and A cells. This is consistent with the observed B1 to B2 conversion described above with the VCT labeling of ventricle contacting cells (Figure 3A, C) and suggests that B2 cells are obligatory relay in the generation of new neurons in the V-SVZ. Analysis of the trajectories starting on B2 cells also included C and A cells (Figure 5F, G). Both B1 and B2 cells showed destination points in B cells (26% for B1; 15% for B2) consistent with a fraction of these cells that self-renew symmetrically^19^ (Figure 5G). This analysis suggests that B1 cell lineages transition through B2 cells and that B2 cells retain self-renewal potential. Importantly, the velocity analysis indicates that B2 cells are progenitors of C cells and of new neurons (A cells).

Interestingly, the majority of active B cells (1473/1816 cells) had a transcriptional profile of B2 cells. From all B2 cells, we found that 43.38% are transcriptionally in an active state. The expression of EGFR is associated with NSC activation and their entry into the cell cycle ^29,55^. To confirm that a fraction of B2 cells is activated *in vivo*, we analyzed their EGFR expression using immunocytochemistry. Immunogold staining under TEM revealed that a fraction of B2 cells, identified by the basal location of their primary cilium-basal body, were EGFR+ (Figure S6L). This was also confirmed with confocal microscopy of coronal sections from hGFAP::GFP mice (Figure 5H). We found that 15.32% ± 4.13 (1,403 cells, n=3) of all B2 cells and 13.5% ±1.62 of the wedge B2 cells were EGFR^+^ (Figure 5I). Previous work has suggested that some B2 cells are actively dividing, but this assessment was not based on the localization of the primary cilium ^22^. We therefore used [^3^H]-thymidine and TEM, and BrdU administration and confocal microscopy, to determine if bonafide B2 cells, identified based on the location of their cilia-basal body, can divide in adult mice. P60 mice received four [^3^H]-thymidine injections, every 2 hrs, and their V-SVZ was analyzed 2 hrs after the last injection. Thirty B cells labeled with [^3^H]-thymidine were reconstructed at the TEM. The primary cilium-basal body revealed that 10 corresponded to B1 (10/30) and 20 to B2 cells (20/30) (Figure S6N-P). This indicates that a fraction of B2 cells divide. To analyze a larger number of B2 cells at a shorter survival after DNA labeling, we injected hGFAP::GFP mice with a single pulse of BrdU and analyzed the V-SVZ 30 min after injection. We found that 2.51% ± 1.43 of B2 cells (5,060 cells, n=3) were BrdU^+^. Proliferative B2 cells were observed in all V-SVZ regions, including the wedge (Wedge: 2.00% ± 0.99, LW-Dorsal: 1.89% ± 0.77, LW-Ventral: 3.18% ± 1.49) (Figure 5J, L). As further confirmation that a fraction of B2 cells is actively dividing, we used the proliferation marker Ki67 in combination with primary cilium-basal body localization in P60 mice. Consistent with the BrdU observations, we found that 2.92% ± 1.05 of B2 cells in the V-SVZ (2,789 cells, n=3) were Ki67+(Figure 5K, L). The above results show that a subpopulation of B2 cells is activated and proliferates *in vivo*.

### B2 cells function as progenitors for adult neurogenesis

The above observations demonstrate that a subpopulation of B2 cells divides. In addition, our transcriptomic analysis suggested that B2 cells give rise to C and A cells in the adult V-SVZ. To directly test if B2 cells can give rise to neurons or glial cells *in vivo,* we took advantage of the wedge region, which contains B2 cells, but lacks B1 cells. We first confirmed that B2 cells in the wedge region do not contact the lateral ventricle. We labeled apical cells by intraventricular injection of VCT into the lateral ventricles of hGFAP::GFP mice (Figure S7A). Twenty-four hours after injection of VCT, ependymal cells and 37.01% ±14.10 of B1 cells were labeled, but B2 cells in wedge were not labeled (n=3) (Figure S7 B-D). Furthermore, B2 cells in the wedge remain VCT negative 3, 7 or 15 days after injection. Isolation of cells from wedge microdissected 24 hours after intraventricular injections of VCT did not contain VCT+ cells (Figure S7E).

We, therefore, injected VCT into the lateral ventricles of mice (P60-P100) carrying constitutive EGFP expression and conditional Ai14. These mice also carried the hGFAP::Cre^ERT2^ transgene to lineage trace hGFAP-expressing cells after transplantation and tamoxifen administration. We microdissected the wedge region from the lateral wall (LW) of the lateral ventricles and separately transplanted dissociated cells from these two regions into the dorsal V-SVZ of P4-5 wild-type C57/Blk6 mice. To induce the translocation of the Cre^ERT2^ (hGFAP::Cre^ERT2^ allele) and expression of TdTomato (Ai14), we administered a single dose of tamoxifen (100mg/kg) immediately after transplantation. This allowed us to lineage trace transplanted B cells in the host brain. The olfactory bulb (OB) and the V-SVZ were analyzed for EGFP^+^/TdTomato^+^ cells 30 days after transplantation (Figure 6A). As expected, Tdtomato^+^/EGFP^+^ granular cells (GCs), and a small number of periglomerular cells (PGCs), were observed in the OB of transplanted mice that received transplants of the LW (Figure S7H-J). Similarly, transplants that included B2 cells from the wedge regions generated Tdtomato^+^/EGFP^+^ granular cells and a small number of EGFP^+^/TdTomato^+^ PGCs (n=6, 3 independent experiments, Figure 6B-D). The GCs derived from wedge B2 cells were primarily localized in the superficial and intermediate granular layers. In contrast, lateral wall transplants that included more ventral V-SVZ regions generated GCs in deeper layers (intermediate and deep layers (n=2; Figure S7I)). The above is consistent with previous findings showing that dorsal and ventral NSCs in the V-SVZ give rise to superficial and deep GCs respectively ^56^. Interestingly, in all mice transplanted with wedge, or with lateral wall cells, similar numbers of Tdtomato^+^/GFP^+^ oligodendrocytes were observed in the corpus callosum (wedge n=6, LW n=2, Figures 6E, F and S7K, L). Mice transplanted with wedge B2 cells, as well as animals that received B cells from the LW, had Tdtomato^+^ cells with migratory morphology in the RMS suggesting that the transplanted cells continue generating progeny one month after transplantation. Together, these results indicate that B2 cells can generate both neurons and oligodendrocytes.

**Figure 6.**
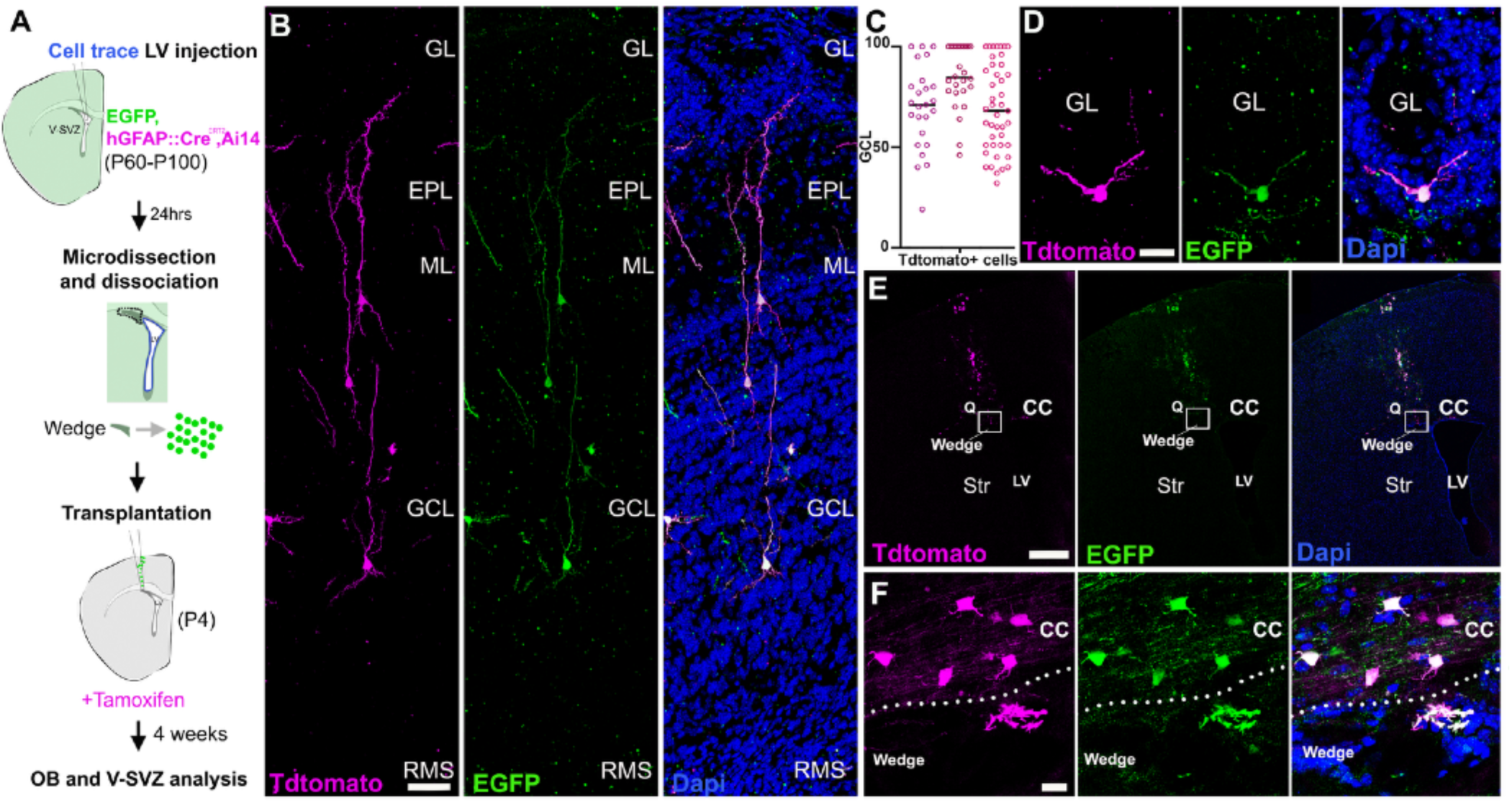
B2 cells are neurogenic. (**A**) Schematic of the wedge transplantation protocol. (**B**) Confocal images of Tdtomato^+^/GFP^+^ granular cells in the OB of a transplanted mouse. (**C**) Granule cell layer (GCL) position of wedge transplant-derived OB granular cell interneurons per animal (symbols represent individual cells). 100 represents superficial GCL. (**D**) Confocal images of periglomerular Tdtomato^+^/GFP^+^ cells. (**E, F**) Transplantation site showing Tdtomato^+^/ GFP^+^ oligodendrocytes in the corpus callosum. GL: Granular layer, EPL: external plexiform layer, ML: mitral layer, GCL: granular cell layer, RMS: rostral migratory stream, CC: corpus callosum, Str: striatum, LV: lateral ventricle. Scale bars: 0.5mm (P), 50μm (M), 35μm (O), 15μm (Q), 10μm (E), 5μm (A, C, H, K), 2μm (B), 500 nm (F).

## Discussion

Here we have addressed how the V-SVZ niche maintains neurogenesis as B1 cells become largely depleted in young adults. Our results indicate that the adult V-SVZ contains NSCs in both the ventricular (B1 cells) and subventricular zone (B2 cells). Our findings suggest that this progenitor function is progressively relayed from B1 to B2 cells during juvenile and early adult stages.

We show that B1 and B2 cells share many structural and gene expression properties, and as previously described, these cells share similarities with parenchymal astrocytes. Therefore, whether B2 cells function as progenitor cells, or instead have a structural role similar to parenchymal astrocytes, remains controversial ^34^. The identification of B2 from B1 cells and parenchymal astrocytes is hindered by the lack of markers that distinguish them. Here, we show that B1 and B2 cells can be identified from parenchymal astrocytes by their location within the V-SVZ and expression of S100a6 ^41^. Moreover, we provide a strategy to distinguish B2 cells from B1 cells by the localization of their primary cilium and basal body. Using this criterion, we found that B2 cells are distributed along the V-SVZ, extending into the dorsolateral expansion of the SVZ, or wedge region ^23,57,58^. B1 cells sharply decreased in number during the first months of life consistent with previous studies ^19,59^. In contrast, B2 cells significantly increased in the juvenile, and remained largely stable until one year of age, after which their number decreased. In a previous study we showed that 20%–30% of B1 cells self-renew symmetrically, and a subpopulation of self-renewing B1 cells loses their apical contact, suggesting conversion into B2 cells ^19^. Here, we used Violet Cell Trace to follow the conversion of B1 into B2 cells and show that this conversion is frequently associated with cell division (Figure 3). Interestingly, velocity analysis suggests that B1 cells may transition through B2 cells in the generation of a neurogenic lineage. However, whether the B2 stage is an obligatory step in the generation of a neurogenic lineage needs to be experimentally confirmed. In addition to the B1-B2 conversion, the transcriptomic analysis suggests that B2 cells can also self-renew, likely contributing to the long-term persistence of B2 cells and neurogenesis.

As indicated above, B1 and B2 cells share ultrastructural, transcriptional, and molecular properties. However, unlike B2 cells, B1 cells are an integral part of the ventricular epithelium. Unsupervised clustering of our single-cell data did not immediately reveal apical/non-apical differences and instead showed two major axes of transcriptional variation among B cells: activated state and regional identity. This observation further illustrates the similarities between B1 and B2 cells and is consistent with previous transcriptomic analyses of the V-SVZ^23–28,31,32,34,43,47^. Selective labeling of B1 cells by adenoviral intraventricular injection, together with microdissections of the wedge region, which contains B2, but not B1 cells, allowed us to identify sets of genes that identify prototypical B1 and B2 cells. Consistent with the B1 cells’ contact with the ventricle ^4^, we found DE genes in B1 cells linked with their epithelial identity (e.g: *Atf3, Actg1, Emp1, Anxa2* ^50,51,60^). These genes were absent or lowly expressed in B2 cells. In contrast, DE genes in B2 cells were frequently associated with the extracellular matrix components (e.g: *Ptprz1, Ptn, Lama2* ^36,61^). These genes were upregulated in B2 cells but also expressed at lower levels in B1 cells. These results suggest the transition between B1 and B2 cells requires the loss of epithelium-associated genes. Therefore, B1 and B2 cells are highly related cell types that form part of a continuum in the populations of astroglial cells within the V-SVZ. Consistently, we identified a population of B cells with transcriptomically intermediate properties between B1 and B2 cells (Figure 4). Given this continuum and the similarities between B1 and B2 cells, the location of the primary cilium-basal body becomes key in defining when B cells are no longer part of the epithelium and have become B2 cells. Future studies using spatial transcriptomics may allow us to determine at what stage, in the transcriptomic transition between B1 and B2 cells, the loss of apical contact occurs. Our protein association network analysis reveals sets of genes coding for interconnected proteins in B1 cells that are associated with epithelial identity. Among these networks, we found genes associated with the transcription factor complex AP-1 (*Atf3, Jun, Junb, Jund*). Notably, this complex is downregulated by *Zeb1*^62^, a transcription factor differentially expressed in B2 cells. Interestingly, *Zeb1* has been linked to delamination from the ventricular cavities ^63,64^ and therefore could be a key player in the transition from B1 to B2 cells.

Our single-cell analysis also shows that B1 and B2 cells can be in quiescent and activated states, suggesting that both populations act as NSCs in the V-SVZ. We found that ∼15% of B2 cells at P60 expressed EGFR, similar to the 11-13% described in B1 cells ^29,35^. BrdU- and Ki67-labeling showed that 2.5% of B2 cells divide, consistent with initial observations using ultrastructure and ^3^[H]-Thymidine labeling ^21^ and also similar to the percentage of all B cells labeled with EdU with 30min survival ^40^. Together, with our transplantation experiment demonstrating that B2 cells generate OB neurons and oligodendrocytes *in vivo*, the above suggests that B2 cells divide, self-renew, and generate progeny. This suggests that B2 cells possess properties of NSCs to sustain neurogenesis in the adult mouse brain. This conclusion is consistent with a recent publication suggesting that basal SVZ cells can function as NSCs in the postnatal brain ^33^. However, the criteria used to identify the basal NSCs in Baur et al. is not consistent with our findings (see results), making a direct comparison between the two studies diffcult.

B2 cells lack direct contact with the CSF, which has been suggested to contain factors that regulate V-SVZ NSCs^17,18,65^. B1 and B2 cells are interconnected via GAP junctions and generate bidirectional calcium waves that propagate along blood vessels^66^. It is possible that through these interconnections, or through ependymal cells, B2 cells derive regulatory information from signals present within the CSF. In the present study, we illustrate how B2 cells are closely associated with blood vessels and the extracellular lamina matrix (ECM) known as fractones^37^. *Lama2*, a protein associated with fractones and blood vessels, was a top DE gene in B2 cells (Table1). Increasing evidence indicates that as the animal ages, more basal signaling from the vasculature and extracellular matrix within the SVZ niche regulates the quiescence of primary progenitors^17,67–69^. Interestingly, fractones increase in size with age ^36,70,71^ and are involved in the regulation of NSCs proliferation^36,72^. In older adults, most of the regulatory mechanisms of NSCs may be linked to increased signaling from ECM and blood vessels.

NSC functions were classically linked to cells within brain epithelia and the VZ ^2,3,73^. More recent studies, primarily in primates, have revealed a large population of basal NSCs in the outer SVZ. These NSCs, identified as outer (basal) RG are considered key to cortical expansion and human brain evolution ^52,74–76^. Here we show, in mice too, a population of non-apical NSCs is essential to maintain long-term neurogenesis in adults. Interestingly, several genes associated with oRG, like *Ptprz1, Ptn, Tnc, Hopx* (Table 1), are also present in B2 cells possibly suggesting shared mechanisms to retain NSC function in cells that are no longer part of the VZ. Non-epithelial NSCs have also been described in the human ganglionic eminences ^77,78^ and are likely essential in the amplification of cortical interneuron production. B1-like cells in the walls of the human lateral ventricles decrease sharply in early postnatal life ^20^. As B1 cells are depleted in non-human primates and humans, a layer of basal SVZ astrocytes (ribbon astrocytes) forms and these cells persist into adults ^79–84^. Whether these cells are derived from B1-like cells and retain the competence to function as primary progenitors for the generation of new neurons or glial cells remains unknown. This basal astroglial cell could be an important reservoir of NSCs for brain repair.

## Supporting information

Table S1_Resources Table

Table 1_DEG_B1_Vs_B2

## Acknowledgments

We thank Dr. McGinnis and Dr. Zev Gartner for the MULTI-seq barcodes and technical advice. We would like to thank members of the Alvarez-Buylla lab for helpful discussions. Work in the Alvarez-Buylla laboratory is supported by Program for Breakthrough Biomedical Research, which is partially funded by the Sandler Foundation, NIH grants P01 NS083513, R01 NS028478, R01 NS113910, R01MH122478 and a generous gift from the John G. Bowes Research Fund. ACS was supported by the Spanish Generalitat Valenciana and European Social Fund (APOSTD2018/A113). JMGV was supported by the Valencian Council for Innovation, Universities, Science and Digital Society (PROMETEO/2019/075).

## Author Contribution

Conceptualization, A.C.-S. and A.A.-B; Methodology, A.C.-S., M.A.N, W.M, S.G.-G, D.M.S and R.R.-R; Formal Analysis, A.C.-S. and M.A.N; Investigation, A.C.-S. and M.A.N; Software, A.C.S and M.A.N; Resources, J.-M.G.-V., D.A.L and A.A.-B.; Writing – Original Draft, A.C.-S., and A.A.-B.; Visualization, A.C.-S; Supervision, K.O, J.-M.G.-V., D.A.L and A.A.-B.; Funding Acquisition, A.C.-S and A.A.-B.

## Supplemental Information

Document S1. Figures S1-S7

Table S1. Resources Table

## Methods

### Mice

Mice were housed on a 12 hr day-night cycle with free access to water and food in a specific pathogen-free facility in social cages (up to 5 mice/cage) and treated according to the guidelines from the UCSF. Institutional Animal Care and Use Committee (IACUC) and NIH. All mice used in this study were healthy and immuno-competent and did not undergo previous procedures unrelated to the experiment. CD1-elite mice, C57BL/6 J, hGFAP::GFP, β-actin-EGFP, hGFAP:CreERt2 and Ai14 mouse lines (see Table S1) were used. Sample sizes were chosen to generate enough high-quality single cells for RNA sequencing, including variables such as sex, and identifying potential batch effects. Biological and technical replicates for each experiment are described in the relevant subsections below.

### Tissue Preparation

Mice were deeply anesthetized with 2.5% Avertin (i.p.) and perfused transcardially with saline (0.9% NaCl), followed by 4% paraformaldehyde (PFA) for immunohistochemistry, 4% PFA -0.5% glutaraldehyde for immunogold staining, or 2% PFA -2.5% glutaraldehyde for transmission electron microscopy. Brains were removed and postfixed in fresh fixative solution overnight, rinsed in 0.1 M phosphate buffer (PB). After cryoprotection in 30% <u>sucrose</u>, fixed tissue was cut on a sliding microtome (Leica SM 2010R) into 30 μm coronal sections and processed as floating sections for immunohistochemistry. For whole mount preparations, brains were removed, and the lateral walls were immediately dissected out as previously described ^85^. Whole mounts were fixed in 4% PFA at 4°C overnight and then stored in PB containing 0.1% sodium azide at 4 C until immunohistochemical processing.

### Lateral ventricle injections

<u>Adenoviral injections:</u> Homozygous (P27-30) Ai14 mice were anesthetized with 250 mg/kg tribromoethanol (avertin) until pedal reflex was abolished and placed in ear bars on a customized stereotaxic rig. After making a small incision, a hole was drilled in the skull at the injection coordinate to expose the brain surface. The brain was then injected with 5ul of adenovirus (Ad:-Cre-GFP) from a beveled pulled glass micropipette. Injection coordinates were determined from a stereotaxic mouse brain atlas and confirmed with dye injections. The x and y coordinates were zeroed at bregma, and the z coordinate was zeroed at the brain surface. Injections at the following coordinates: L:0.8, A:0.17, D: 2.25. The incision was closed with silk suture, and animals were placed on a warm surface and monitored until they resumed feeding and grooming activity. Animals were used between 20-24 hrs after injections for microdissections and scRNA-sequencing analysis. <u>Violet Cell trace injections:</u> P14 hGFAP:GFP animals were anesthetized and received an intraventricular injection of 3ul of Violet Cell trace 2.5mM in saline.

### [3H] Thymidine and BrdU administration

To identify proliferating cells by TEM, we administered four intraperitoneal injections of 3H-Thy at 2 hrs intervals to adult mice (1.67 μL/g body weight, specific activity 5 Ci/mmol; PerkinElmer) with subsequent perfusion 2 hrs after the last injection (n = 3). Brdu: To detect proliferating cells we injected a single dose of Brdu (50mg/kg) in sterile saline and perfused 30 min after the injection. To detect label retaining cells, BrdU was dissolved in water at 1 mg/ml and administered in the drinking water.

### Immunohistochemistry

Whole-mounts were incubated in primary and secondary antibodies (see Table S1) in PBS with 0.2% TX-100 and 10% normal goat serum for 48 hrs at 4°C. For samples processed for BrdU unmasking, incubations with primary and secondary antibodies were performed first, followed by fixation in 4% PFA at room temperature for 10 minutes. Tissue was then incubated in 2N HCl at 37°C for 30 minutes, followed by treatment with 0.1M boric acid for 20 minutes at RT prior to incubation with anti-BrdU antibodies and corresponding secondary antibodies for 48 hrs at 4°C each. Coronal sections were processed according to the same protocols but with shorter antibody <u>incubation times</u> (primary antibodies: overnight at 4°C; secondary antibodies: 2 hours at room temperature). Primary antibodies are listed on the methods table. Secondary antibodies were conjugated to Donkey AlexaFluor dyes (Invitrogen/Molecular Probes).

### Transmission electron microscopy

For TEM, mice were fixed with 2% PFA-2.5% glutaraldehyde. Brains were rinsed in 0.1 M phosphate buffer (PB) and cut into 200-mm sections. Sections were post-fixed in 2% osmium tetroxide, dehydrated, and embedded in Durcupan resin (Fluka; Sigma-Aldrich). Semithin sections (1.5 mm) were cut with a diamond knife and stained with 1% toluidine blue for light microscopy. Ultrathin sections (70–80 nm) were cut, stained with lead citrate, and examined under an FEI Tecnai G2 Spirit transmission electron microscope (FEI Europe) using a digital camera (Xarosa (20 Megapixel resolution), Radius EMSIS GmbH, Münster, Germany). Immunogold staining and TEM processing was performed as previously described ^86^.

### TEM serial reconstructions

For B1 and B2 cells serial reconstructions, 150 serial ultrathin sections (70nm) from the V-SVZ (ages: P10, P60 and P365, n=1) were imaged. Each B cell within the V-SVZ was followed along all ultrathin sections. Ultrastructural features of each cell were annotated. B1 and B2 cell identities were assigned based on the apical or basal localization of their primary cilium-basal body. Full reconstructions confirmed that B2 cells do not show apical contact. Cells that were not fully reconstructed were not included in the study. For 3D reconstruction on Figure 1 serial TEM micrographs were aligned with FIJI TrakEM2 software and rendered using the surface tool manually on Imaris.

### Autoradiography

Brains injected with 3H-Thy were processed for TEM as described above. Subsequently, V-SVZ semithin sections were dipped in autoradiography emulsion (Carestream Autoradiography Emulsion, Type NTB), dried in the dark, and stored at 4°C for 4 weeks ^35^. Autoradiography was developed using standard methods and counterstained with 1% toluidine blue. Close to the LV 3H-Thy-labeled nuclei were identified in semithin sections. Six or more silver grains needed to be present over the nucleus, and the nucleus had to be labeled in at least three consecutive serial sections, for a cell to be considered labeled. All consecutive sections showing labeled cells were selected under a light microscope (Eclipse; Nikon). Consecutive sections showing labeled cells were selected under a light microscope (Nikon, Eclipse), re-embedded, and ultrathin-sectioned (70nm) for <u>TEM</u> analysis (Tecnai Spirit G2, FEI, Oregon). Anatomical references were used to identify labeled cells under EM.

### B1 and B2 cell maps

<u>Coronal Section Maps</u>: Serial coronal sections were selected (Bregma 1.94, 1.70, 1.42, 1.10, 0.86, 0.14, -0.10) and immunostained for GFP, Arl13b, γ-tubulin and S100a6 (n=2, P30). Confocal Z-stacks (Z size=20μm, Z step size=0.5) of the full length of the V-SVZ/section were obtained using an oil 63x objective. Images were stitched and analyzed using Imaris software 3D view. We used the ortho-slicer function and spots tool to move across planes and manually add B1 and B2 spots. The cytoplasm of B1 and B2 cells was identified by GFP and S100a6 expression within the V-SVZ. B1 or B2 cell labels were assigned based on the apical or basal location of their primary cilium-basal body (Arl13b and γ-tubulin). Spots were always placed on the cell basal body. Spot maps were generated with Imaris, organized and annotated with Adobe Photoshop. <u>Whole mount map</u>: A whole mount (P60, female, n=1) was dissected, fixed, and stained for ß-Catenin, GFP, Arl13b and γ-tubulin. Confocal Z-stacks of 129 V-SVZ positions were obtained with a 63x oil objective. The beginning of z-stacks were set on the multiciliated ependymal cells cilia tips (identified by Arl13b) to ensure that all apical cells were included in the analysis. The end of each stack was set deep into the tissue to include the whole thickness of the SVZ. We considered that we reached the striatum when the DAPI and GFP^+^ cells were no longer aggregated in cell clusters (Z size was variable between regions, Z step size=0.5). Images were stitched and analyzed as above. To map the high density of spots, we obtained a csv file with all spots positions in Imaris and we imported this data to Rstudio. We used the ggplot package to visualize B1 and B2 spots.

### Quantifications

<u>Cilium orientation:</u> Coronal sections from hGFAP::GFP mice (P30, n=3, 2 sections/mouse) were selected and immunostained for GFP, Arl13b, γ-tubulin and S100a6. Confocal Z-stacks (Z size=20μm, Z step size=0.5) of the full length of the V-SVZ/section were obtained using a63x oil objective. Images were stitched and analyzed using Imaris software 3D view. To determine cilium orientation, a transparent radar diagram was superimposed on the images. The ventricular zone, defined by multiciliated ependymal cells contacting the ventricular cavity, was considered 0 degrees and the striatum 180 degrees, dorsal: 45, 90, 145 degrees and ventral: -45, -90 and -145 degrees. Spots were manually added to the primary cilium of B2 cells. V-SVZ dorsal and ventral regions were analyzed. Radar charts were obtained with Excel. <u>B1-B2 cells</u> <u>over time:</u> Whole-mounts from hGFAP:GFP mice (P10, P21, P28, P60, P90, P120, P180 and P365 (n= 3/age/sex) and P545 (3 females, 1 male)) were dissected, fixed, and immunostained for GFP, ß-Catenin, γ-tubulin and Arl13B. Confocal Z-stacks of 9 positions/region (antero- ventral and posterior-dorsal)/mouse were obtained using a 63x oil objective. The beginning and end of Z-stacks, B1 and B2 cells’ identification and analysis was performed as above. <u>Violet</u> <u>Cell trace quantifications:</u> hGFAP:GFP mice injected with VCT at P14 were analyzed at 1, 15 and 30 days after injection. Coronal sections (30μm) from the contralateral hemisphere were stained for GFP, γ-tubulin and Arl13B or GFP, γ-tubulin, Arl13B and BrdU. Images from the wedge or lateral wall (including dorsal and ventral regions) were obtained using a 63x oil objective (Z size=30μm, Z step size=0.5). For B1-B2 cell conversion: 8 sections/mouse, n=3, d1: 5277 cells analyzed, d15: 4456 cells analyzed, d30: 4008 cells analyzed. For B2 cells, Brdu^+^VCT^+^: 6 sections/mouse, n=4, 284 Brdu^+^ LRCs analyzed. <u>Adenoviral infection quantifications:</u> Whole mounts from the contralateral hemisphere from Tdtomato mice (P28, n= 3) were dissected, fixed, and immunostained for Tdtomato, ß-Catenin, γ-tubulin and Arl13B. Note that only the contralateral hemisphere of the injection was used for analysis. Confocal Z-stacks of 6 positions/region (antero-ventral and posterior-dorsal)/mouse were obtained at 63x, as above. 1995 B1 cells and 4446 ependymal cells were analyzed. <u>EGFR+, Brdu+ or Ki67 B2</u> <u>cells:</u> Quantifications of EGFR, Brdu or Ki67 were performed on coronal sections from P30 hGFAP:GFP mice co-stained with Arl13b and γ-tubulin. Images from the wedge, dorsal, and ventral lateral wall were obtained using a 63x oil objective (Z size=30μm, Z step size=0.5). B2 cells were identified as above (EGFR: 6 sections/mouse, n=3, 1403 B2 cells analyzed. Brdu: 12 sections/mouse, n=3, 5060 B2 cells analyzed. Ki67: 6 sections/mouse, n=3, 2789 B2 cells analyzed). <u>GCL position:</u> Confocal 10x images were obtained from non-consecutive OB sections (825 cells analyzed, n=6). Relative positions of Tdtomato^+^ cells within the GCL were calculated with the Fiji Straight tool. The position of Tdtomato^+^ cells was calculated as a percentage of GCL thickness (0=Deep, 100=Superficial).

### Statistics

All results shown in the graphs are expressed as mean ± SD. Statistical significance was defined as * p ≤ 0.05, ** p < 0.01, *** p < 0.001, **** p < 0.0001. One-way ANOVA with Tukey’s multiple comparisons (Prism GraphPad software) was used to determine statistical significance of multiple comparisons; Student’s t test (Excel) was used for pairwise comparisons between two groups.

### V-SVZ transplantation

To confirm that wedge transplants did not contain apical cells, we injected 1 ul Cell trace, Violet (C34557, ThermoFisher) into the posterior lateral ventricle of EGFP, hGFAP::Cre^ERT2^,Ai14 mice (coordinates, bregma x:-0.45, y: -0.29, z:3). 24 hrs after Cell Trace injection, the V-SVZ dorsal wedge and lateral walls were microdissected in L-15 medium. Tissue was digested for 30 mins at 37°C in a thermomixer at 900 RPM. Cells were centrifuged for 5 min, 300 RCF at room temp, and the pellet was resuspended with a DNAase/ovomucoid inhibitor according to manufacturer’s protocol (Worthington). To remove myelin the cell suspension was incubated with Myelin Removal Beads (130-096-733, Miltenyi Biotec) for 15 mins at 2-8°C. Cells were washed with 0.5% BSA-PBS and transferred to MACS columns (30-042-401 and QuadroMACS Separator 130-090-976, Miltenyi Biotec). Cells were centrifuged and resuspended in L-15 with DNAse I (180 ug/ml). The dissociated cells were then concentrated by centrifugation (4 minutes, 800xg) and cell suspensions were loaded into beveled glass micropipettes (≈70-90 μm diameter, Wiretrol 5 μl, Drummond Scientific Company) prefilled with mineral oil and mounted on a microinjector. Recipient mice (C57Bl/6, P4) were anesthetized by hypothermia (∼4 minutes) and positioned in a clay head mold that stabilizes the skull ^56^. Micropipettes were positioned at an angle of 0 degrees from vertical in a stereotactic injection apparatus. Injections were performed in the left hemisphere. Eye coordinates were x: 1.7, y: 3.2, z:1.3 from the surface of the skin. After injection, mice received an intraperitoneal single dose of tamoxifen 100mg/kg. Mice were returned to their littlers. For hGFAP::Cre^ERT2^,Ai14 leak controls, we did not administer tamoxifen and kept transplanted mice in a different cage. Three weeks after transplantation, Brdu (5mg/ml) was administered in drinking water for 7 days.

### 10x Technology single cell sample preparation and multiplexing

Samples from mice injected with Ad:Cre:GFP were prepared as following: 15-24 hours after Ad:Cre:GFP intraventricular injections, mice (Experiment 1: n=14, P27-30, Experiment 2: n=12, P27-p30) received intraperitoneal administration of 2.5% Avertin followed by decapitation. Brains were extracted and 0.5 mm slices were obtained with a mouse brain slicer (Steel Brain Matrix - Coronal 0.5mm, Alto). The V-SVZ lateral wall and dorsal wedge were microdissected and considered separate samples. V-SVZ micro-dissections were performed in L-15 medium on ice and the tissue was transferred to Papain-EBSS (LK003150, Worthington). Tissue was digested for 30 mins at 37°C in a thermomixer at 900 RPM. Mechanical dissociation with a P1000 pipette tip (20 seconds), then with a fire-polished pasteur pipette for 3 min. Cells were centrifuged for 5 min, 300 RCF at room temp, and the pellet was resuspended with DNAase/ ovomucoid inhibitor according to the manufacturer’s protocol (Worthington). Cells were incubated in red blood cell lysis buffer (420301, Biolegend) 3-4 min at 4°C. For MULTI-seq barcoding, cells were suspended with Anchor:Barcode solution for 5 minutes at 4°C. A Co-Anchor solution was added and incubated for 5 mins ^87^. Samples were combined and filtered with a FlowMi 40µm filter (BAH136800040-50EA, Sigma). To remove myelin, the cell suspension was incubated with Myelin Removal Beads (130-096-733, Miltenyi Biotec) (6μl/ brain) for 15 mins at 2-8°C. Cells were washed with 0.5% BSA-PBS and transferred to MACS columns. The effluent was collected, and the cell density was counted. Cells were loaded into a 10x Genomics Chromium Single Cell Controller. We used the 10x Genomics Chromium Single Cell 3’ Library & Gel Bead Kit v3.1 to generate cDNA libraries for sequencing according to the manufacturer’s protocols. We confirmed that isolated cells from wedge dissections did not contain apical cells (GFP^+^ and/or Tdtomato^+^ cells) under an epifluorescence microscope.

### Sequencing and read alignment

Libraries were sequenced in an Illumina NovaSeq 6000. Alignment was performed on STAR v2.7.11 and included both intronic and exonic mappings (*--soloFeatures Gene GeneFull Velocyto*). We used an optimized mouse reference genome ^88^ edited to include *GFP* and *Cre* genes. We also included a WPRE sequence to serve as a proxy for TdTomato expression (as it is present in the 3’ end of the TdTomato gene in ^47^).

### Single cell RNAseq data normalization and dimensionality reduction

Cell barcodes were filtered to remove barcodes associated with multiple Multi-seq barcodes, high mitochondrial reads (> 10% reads in mitochondrial genes). We also filtered out barcodes in the bottom or top 2% of total number of reads in order to remove low-quality cells and doublets not detected by Multi-seq barcoding, respectively. Data from different technical replicates were integrated for batch correction (Stuart 2019) using CCA integration in Seurat v. 5 ^89^. Counts were normalized using a negative binomial model (SCTransform v2), dimensionality was reduced to 100 principal components (PCs), which were used to project in two dimensions using uniform manifold approximation and projection (UMAP).

### Identification of transcriptional profiles of B1 and B2 cells

After intraventricular injection of Ad:-Cre-GFP virus, cells contacting the cerebrospinal fluid (ependymal and B1 cells) were infected and identified by their expression of *Cre*, *GFP* and *TdTomato.* Since only 25% of B1 cells are infected after intraventricular injection, we identified the 3 nearest neighbors on the gene expression space for each identified B1 cell. We then performed differential expression analysis between B1 cells (labeled and inferred) and B2 cells. We identified the differentially expressed genes in B1 cells and selected genes for validation with RNAscope. The criteria to select genes for validation was based on: 1) exclusion of genes associated with regionality, 2) genes that were expressed in both adenoviral-infected and non-infected datasets, 3) genes showing V-SVZ expression *in situ* hybridization (ISH) data from the Allen Mouse Brain atlas. We scored cells based on the combined expression of these validated B1 genes using the *AddModuleScore*() in Seurat (B1 score).

### Protein-protein interaction networks

We used Cytoscape with STRINGapp to visualize protein to protein interaction networks between differentially expressed genes in B1 cells (804 protein coding genes, adjusted p-val: 0.01) or B2 cells (303 protein coding genes, adjusted p-val: 0.01). A Mus musculus full STRING network with a confidence score of 0.9 was used as reference. First neighbor interactions were selected for *Actg1, Atf3*, *Anxa2* and *Ptprz1*. STRING functional enrichment analysis was performed in each one of the selected networks.

### Lineage trajectory analysis

Lineage trajectory analyses were performed on two sets of data: one containing only B cells and another containing the neurogenic lineage. B cells were selected and reclustered, cells in the neurogenic lineage were selected and reclustered, but the impact of cell cycle in the dimension reduction was mitigated by regressing out the cell cycle scores (G2.score and M.score) calculated in Seurat. Spliced and unspliced count matrices for these cells were generated using the STAR aligner (2.7.11) with *–soloFeatures Gene Velocyto* option on an optimized mouse genome reference^88^. Lineage trajectory analysis was performed using the scvelo package (v. 0.3.1) in python (v. 3.1), *scvelo.pp.moments*() function was executed using 30 neighbors and 30 PCs, *scvelo.tl.velocity*() was executed with default parameters and *scvelo.tl.velocity_graph*() was executed using the UMAP coordinates calculated in the initial reclustering in Seurat as the *basis* argument. Typical trajectories arising from B1 and B2 cells were inferred using simulated cell state transitions along the neurogenic lineage using *scvelo.utils.get_cell_transitions* with 20 neighbors and 300 steps.

### RNAscope assay

For Coronal or *en face* sections, hGFAP::GFP Mouse brains (P30) were serially sectioned using a Leica cryostat (12 um-thick sections in Superfrost Plus slides) (n=2-4 mice/probe). Sections were incubated 10 min with 4% PFA and washed 3×10 min with phosphate-buffered saline (PBS) to remove OCT. Slides were incubated with ACD hydrogen peroxide for 10 min, treated in 1x target retrieval buffer (ACD) for 5 min (at 96–100 °C) and rinsed in water and 100% ethanol. Samples were air dried at 60°C for 15 min and kept at room temperature overnight. The day after, samples were treated with Protease Plus for 15 min at 40 °C in the RNAscope oven. Hybridization of probes (see Table S1) and amplification solutions was performed according to the manufacturer’s instructions. Amplification and detection steps were performed using the RNAscope 2.5 HD Red Detection Kit, RNAscope 2.5 HD Duplex Reagent Kit or Multiplex. DapB mRNA probe (cat.# 310043) was used as negative and Mm-Crym as positive control. RNAscope assay was directly followed by antibody staining for chicken anti-GFP or anti-ß-Catenin and mouse anti-γ-tubulin. Samples were blocked with TNB solution (0.1 M Tris– HCl, pH 7.5, 0.15 M NaCl, 0.5% PerkinElmer TSA blocking reagent) for 30 min and incubated in primary antibodies overnight. Samples were washed with PBS-Tx0.1% and incubated with secondary antibodies Donkey anti-Chicken biotinylated (Jackson ImmunoResearch, 1:500) and Donkey anti-mouse 647 (Jackson ImmunoResearch, 1:500) in TNB buffer for 1.5 hrs. Samples were washed and incubated with Streptavidin HRP (1:200 in TNB solution) for 30 min. Washed 3×5 min and incubated with Fluorescein Tyramide 5 min (1:50 in amplification diluent) rinsed and incubated with DAPI 10 min. Sections were mounted with Prolong glass Antifade Mountant (Invitrogen, P36980).

## Supplementary Figures

**Figure S1.**
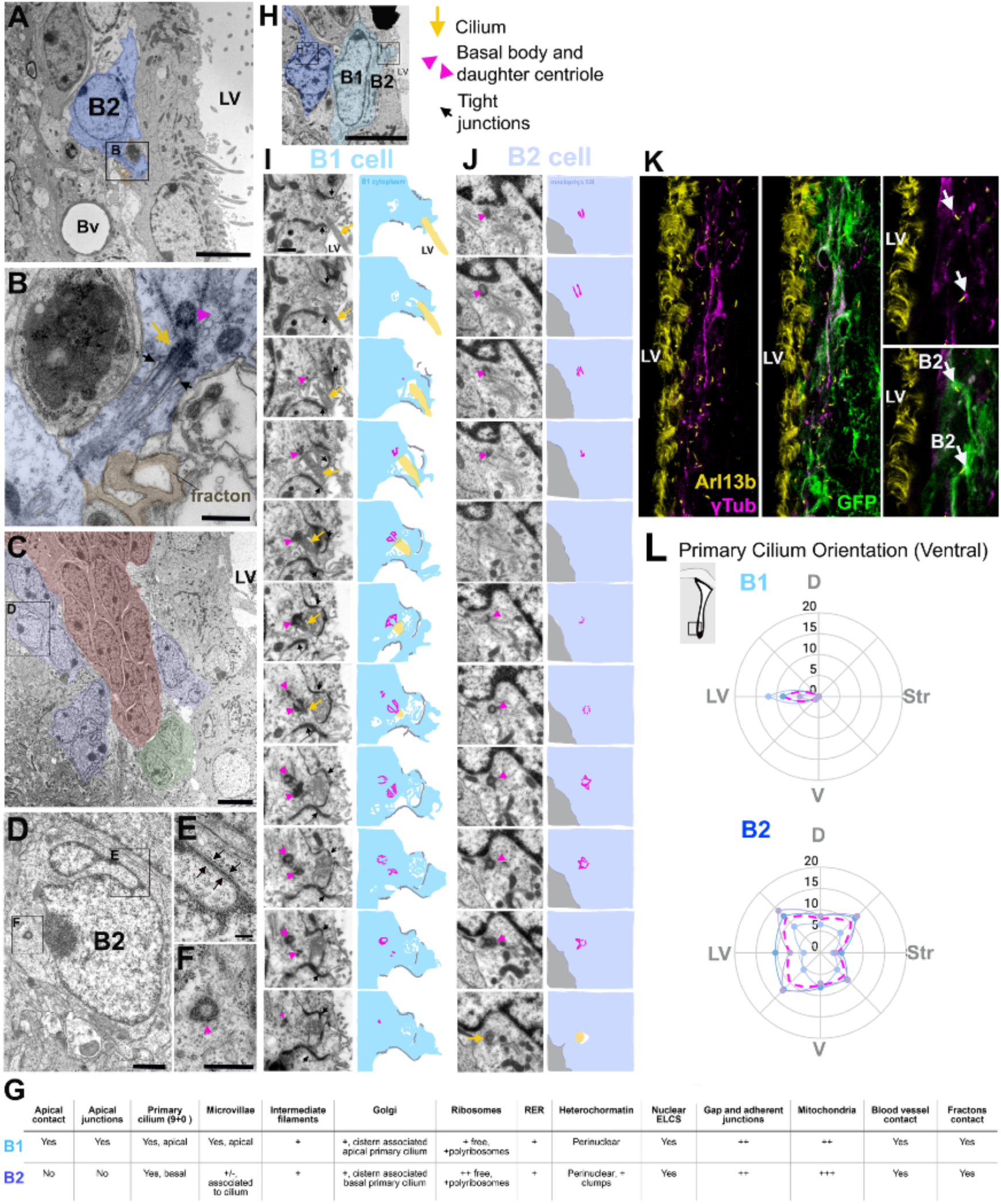
(**A-J**) V-SVZ TEM micrographs. (**A**) B2 cell pseudo-colored in blue. (**B**) High magnification of the B2 cell in (A) contacting fractones (pseudo-colored in orange). This expansion contains a primary cilium (yellow arrow) with ciliary pockets (black arrows) and its daughter centriole (pink arrowhead). (**C**) Image of the V-SVZ B2 cells (pseudo-colored in blue) ensheathing a chain of neuroblasts (A cells, pseudo-colored in red) and intermediate progenitors (C cells, pseudo-colored in green). (**D-F**) Higher magnification of a B2 cell in (C) showing a nuclear envelope chromatin sheet (ELCS) specialization (arrows in E) and its daughter centriole (F). (**G**) Table of B1 and B2 cell ultrastructural characteristics. (**H**) TEM micrograph of B1 (light blue) and B2 (dark blue) cells. (**I**) Serial reconstruction of B1 cell apical contact showing its primary cilium (yellow arrow) and daughter centriole (pink arrowhead). This contact shows tight junctions (black arrows) with neighboring cells. Schematic of B1 cell apical contact (right). (**J**) Serial reconstruction of B2 cell primary cilium (yellow) and its daughter centriole (pink arrowhead). Note the lack of tight junctions. Schematic of B2 cell primary cilium-basal bodies (right). (**K**) Confocal images of a V-SVZ coronal section showing GFP^+^ B2 cells and their primary cilium-basal body labeled with Arl13B (yellow) and γ-tubulin (magenta). (**L**) Radar maps of Ventral B1 and B2 cells’ primary cilium orientation. Average of B2 cells primary cilium localization shows random orientation (magenta, n=3, 394 cells). LV: Lateral Ventricle, Str: Striatum. Scale bars: 5μm (A, C), 500nm (B, D and F), 2.5μm (E) and 2μm (H).

**Figure S2.**
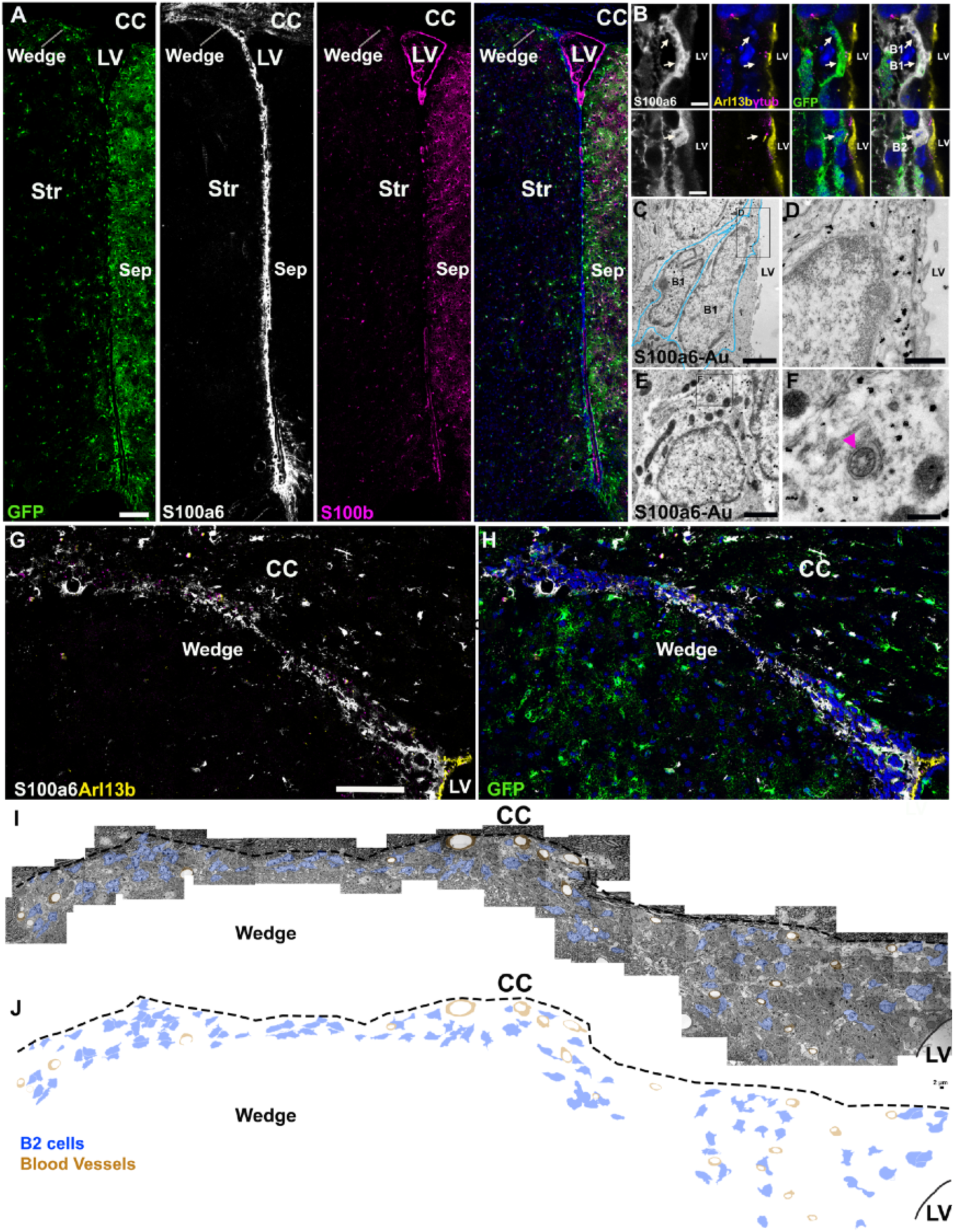
(**A, B**) Confocal images of the V-SVZ from a hGFAP:GFP mouse. High S100a6 expression was found in the V-SVZ. Ependymal cells and parenchymal astrocytes were identified by S100b expression. (**B**) B1 and B2 cells expressing S100a6. (**C-F**) TEM micrographs of S100a6 immunogold staining showing labeled B1 (C, D) and B2 cells (E, F). Cell profile is delineated in B1 (light blue). B2 cell identity was confirmed by the basal location of its primary cilium (pink arrowhead). (**G-H**) Confocal images of the Wedge region from a hGFAP:GFP mouse. Note that the wedge is devoid of ventricular cavities and contains S100a6^+^ B2 cells. (**I-J**) Tiled TEM micrographs showing B2 cell distribution in the wedge (pseudo-colored in blue). LV: lateral ventricle, Str: Striatum, Sep: Septum. Scale bars: 150μm (A), 80μm (H-I), 5μm (B), 2μm (C, I,J), 500nm (D), 2μm (E) and 250 nm (F).

**Figure S3.**
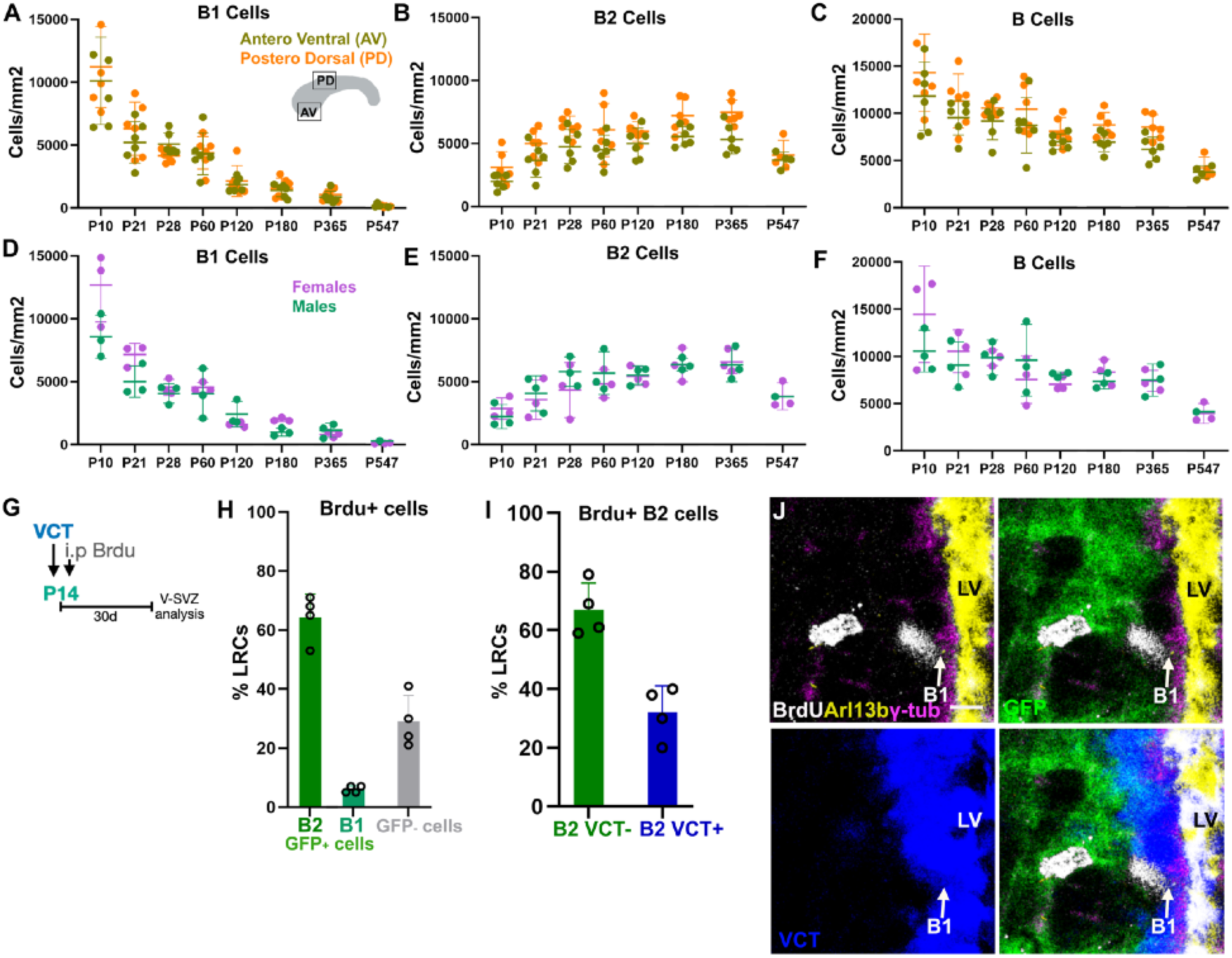
(**A-C**) Quantifications of B1, B2 and B cells over time in the Antero-Ventral and Postero-dorsal V-SVZ domains (n=6/time point, except P547 n=4). No significant differences were observed. (**D-F**) Quantifications of B1, B2 and B cells over time in males and females (n=3/time point). No significant differences were observed. (**G**) Experimental outline: VCT intraventricular injection followed by one pulse of Brdu at P14. The contralateral V-SVZ was analyzed 30 days after injection. (**H**) Percentage of GFP^+^ and GFP^-^ label retaining cells (LRCs). GFP^+^ cells were identified as B1 and B2 cells. (B1 cells: 6.14% ± 1.26, B2 cells: 64.46% ± 8.00 and 29.40% ± 8.99 GFP cells, n=4, 284 cells). (**I**) Out of 181 Brdu^+^ B2 cells, 32.48% were labeled with VCT. Each symbol represents one animal. (**J**) Confocal images of a B1 cell positive for BrdU and VCT, close to a VCT^-^ B2 cell. LV: lateral Ventricle. Scale bars: 5μm (J).

**Figure S4.**
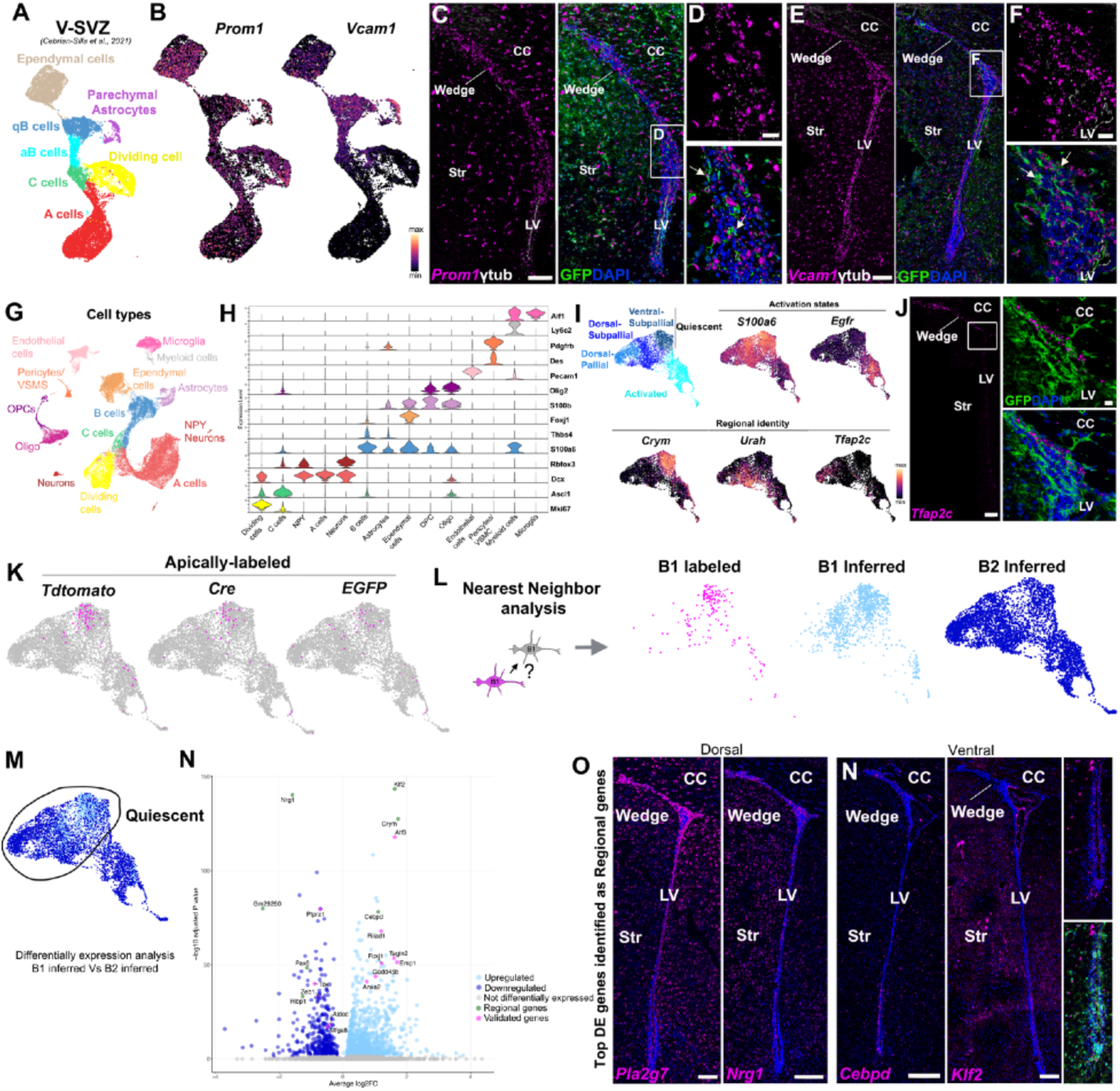
**(A)** UMAP plot of the V-SVZ dataset from ^23^. **(B)** *Prom1* and *Vcam1* expression in the V-SVZ. (**C-D**) Confocal images of *Prom1* RNA (magenta) (C, D) and *Vcam1* RNA (magenta) (E, F) in hGFAP:GFP mice. High magnification images show that *Prom1* and *Vcam1* are expressed in the wedge and in B2 cells (arrows), identified by GFP expression (green) and the localization of basal bodies (γ-tubulin, white). (**G, H**) UMAP plot of scRNA-Seq cell types captured after sequencing and downstream analysis. Violin plot of cell-type-specific marker expression. (**B**) Gene expression identifies B cell clusters’ main sources of heterogeneity: activation states (ativated: *S100a6^low^* and *Egfr+*) and regionality (Pallial: *Tfap2c*^+^, Dorsal-subpallial: *Urah*^+^/Crym^-^ and Ventral-Subpallial: *Urah*^-^/Crym^+^). (**J**) Confocal images of *Tfap2c* RNA (magenta) and GFP protein expression in the V-SVZ. Note its restricted expression in the Pallial V-SVZ domain colocalizing with GFP^+^ cells (B2 cells). (**K**) Distribution of apical-labeled cells (B1 cells) identified by the expression of *Tdtomato, Cre* and *GFP.* (**L**) Distribution of inferred B1 and B2 cells obtained with nearest neighbor analysis. (**M**) Volcano plot of differentially expressed genes between quiescent B1 and B2 cells. (**N**) Confocal images of B1 and B2 differentially expressed genes that show regional distribution. CC: corpus callosum, Str: Striatum, LV: lateral ventricles. Scale bars:100μm (C, E, J, O, N), 20μm (D, F) and 10μm (J).

**Figure S5.**
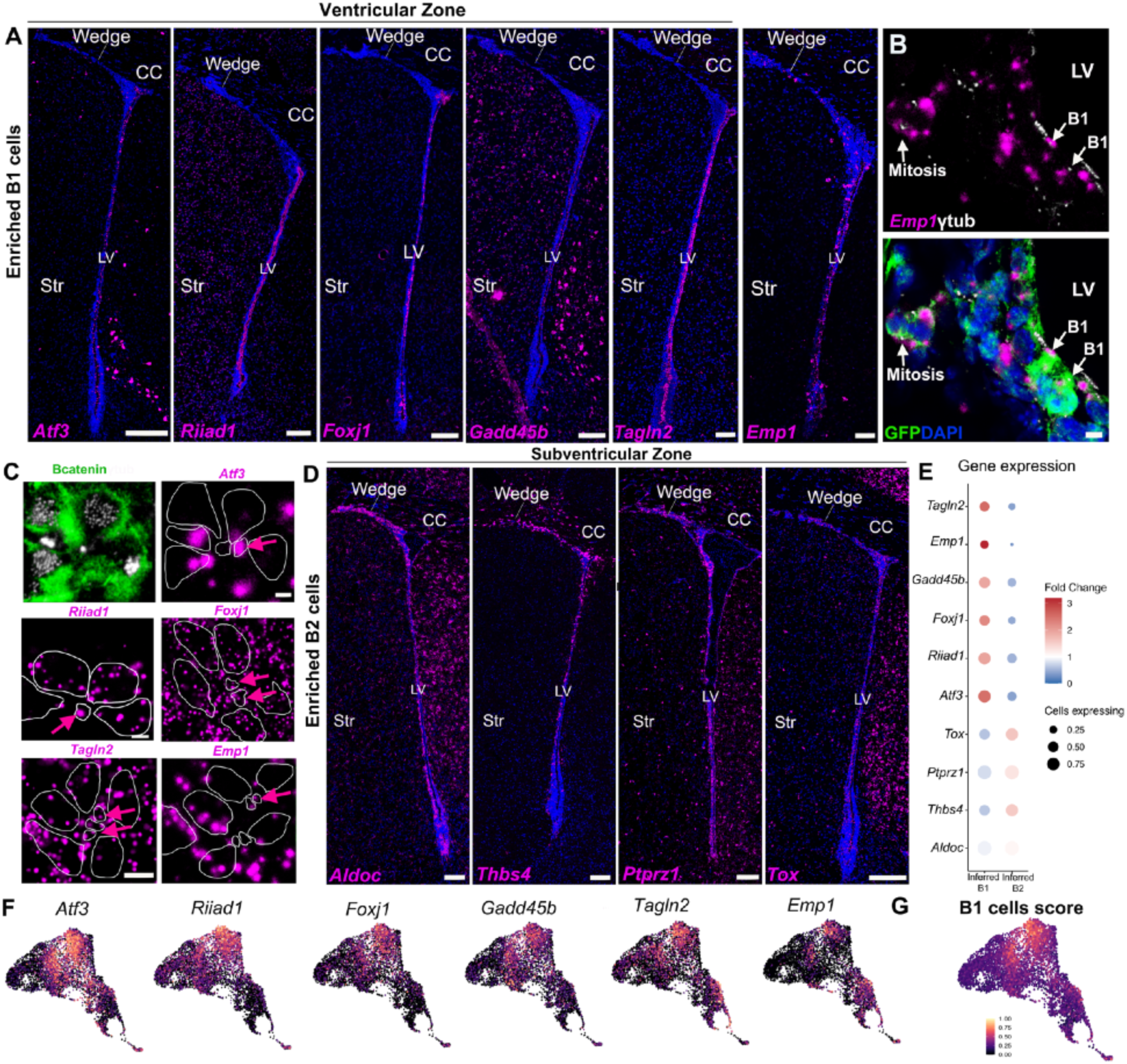
(**A**) Confocal images depict RNA expression of B1 cells differentially expressed genes in the ventricular zone (VZ). (**B**) High magnification confocal images of *Emp1* RNA in combination with GFP and γ-tubulin (white) showing B1 cells and mitotic figures expressing *Emp1.* (**C**) Confocal images of *en face* sections of the ventricular wall reveal the expression of *Atf3*, *Riiad1*, *Foxj1, Tagln2 and Emp1* (magenta) on B1 cells (arrows). B1 cells apical contacts were identified at the center of the pinwheels delimited by ß-Catenin (green) and containing a primary cilium (γ-tubulin, white). ß-Catenin expression is delineated to facilitate RNA visualization. (**D**) Confocal images depict RNA expression of B2 cells differentially expressed genes in the ventricular-subventricular zone (V-SVZ). (**E**) Dotplot showing gene expression of B1 and B2 cells differentially expressed genes in (A-D). (**F**) Gene expression of B1 cell genes. (**F**) Distribution of scored B1 cells. CC: corpus callosum, Str: striatum, LV: lateral ventricles. Scale bars: 100μm (A, D), 5μm (B) 5 and 10μm (C).

**Figure S6.**
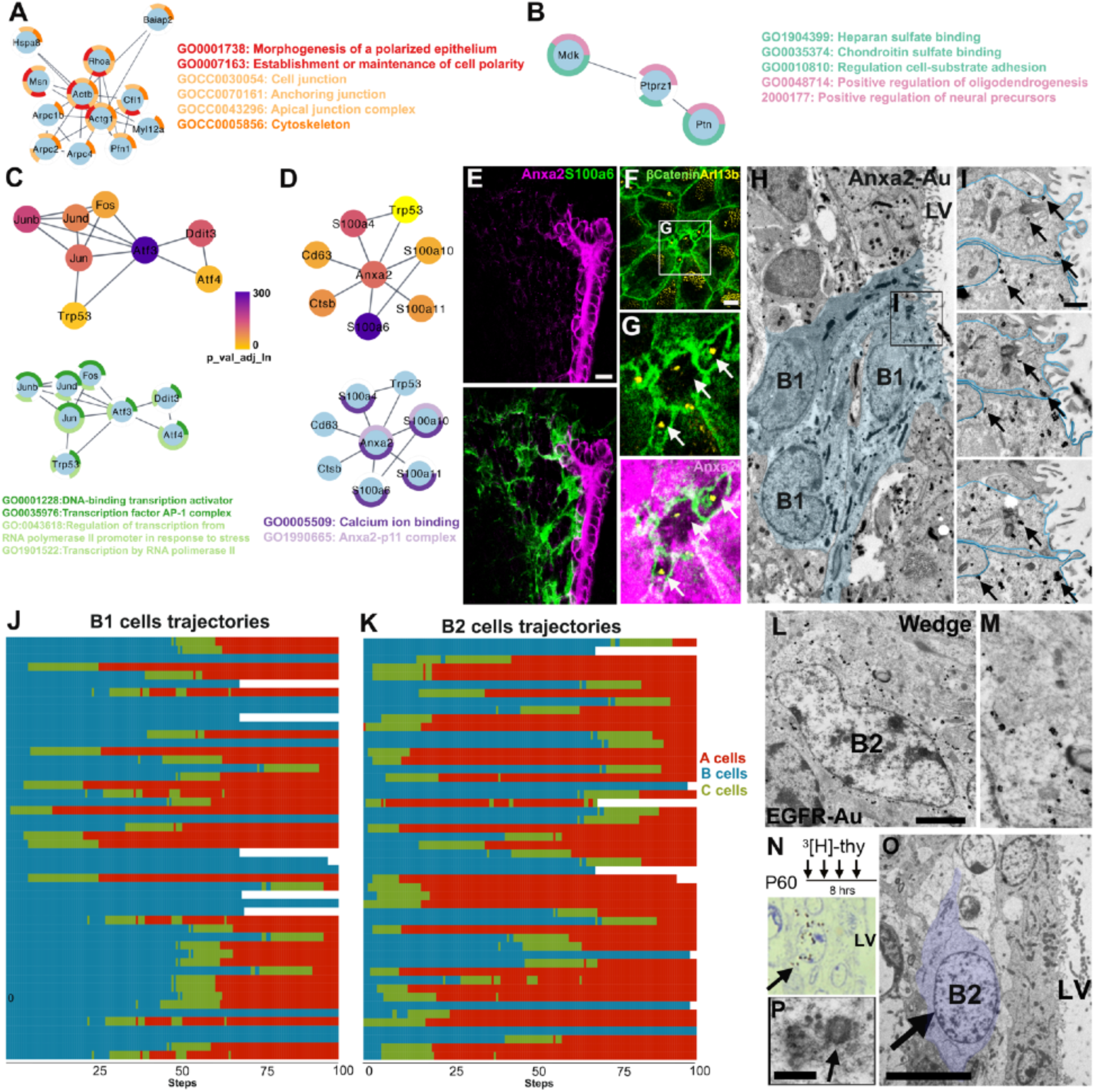
(**A-D**) Protein-protein network diagrams for differentially expressed genes in B1 (A, C, D) and B2 cells (B). (**A**) *Actg1* first neighbors’ network with their functional enrichment annotations. (**B**) *Ptprz1* first neighbors’ network with their functional enrichment annotations. (**C, D**) *Atf3* (C) and *Anxa2* networks with their first neighbors and functional enrichment annotations. (**E**) Confocal images of Anxa2 and S100a6 protein in the dorsal V-SVZ. (**F, G**) Confocal images of whole mount preparation showing B1 cells at the center of the pinwheels expressing Anxa2 on their apical contact (arrows). (**H, I**) TEM micrographs of Anxa2 immunogold staining showing expression in the apical contact of B1 cells (arrows). (**I**) Images of three serial sections from cells in (H). (**J, K**) Barplots showing a sample of 50 B1 and 50 B2 cell trajectories along the neurogenic lineage. **(L, M)** TEM micrographs of an EGFR^+^ B2 cell in the wedge. (**N**) Autoradiography of the V-SVZ of a P60 mouse after receiving 4 ^3^H-Thymidine (^3^H-Thy) injections. The labeled cell was identified on toluidine blue-stained semithin sections (arrow). (**O, P**) TEM micrographs of the ^3^H-Thy-labeled cell in (N), identified as a B2 cell with light cytoplasm, irregular countourn, lack of apical contact, and basal localization of the basal body (arrow) (P). Scale bars: 50μm (E), 2μm (F), 1μm (L) and 500nm (I, P).

**Figure S7.**
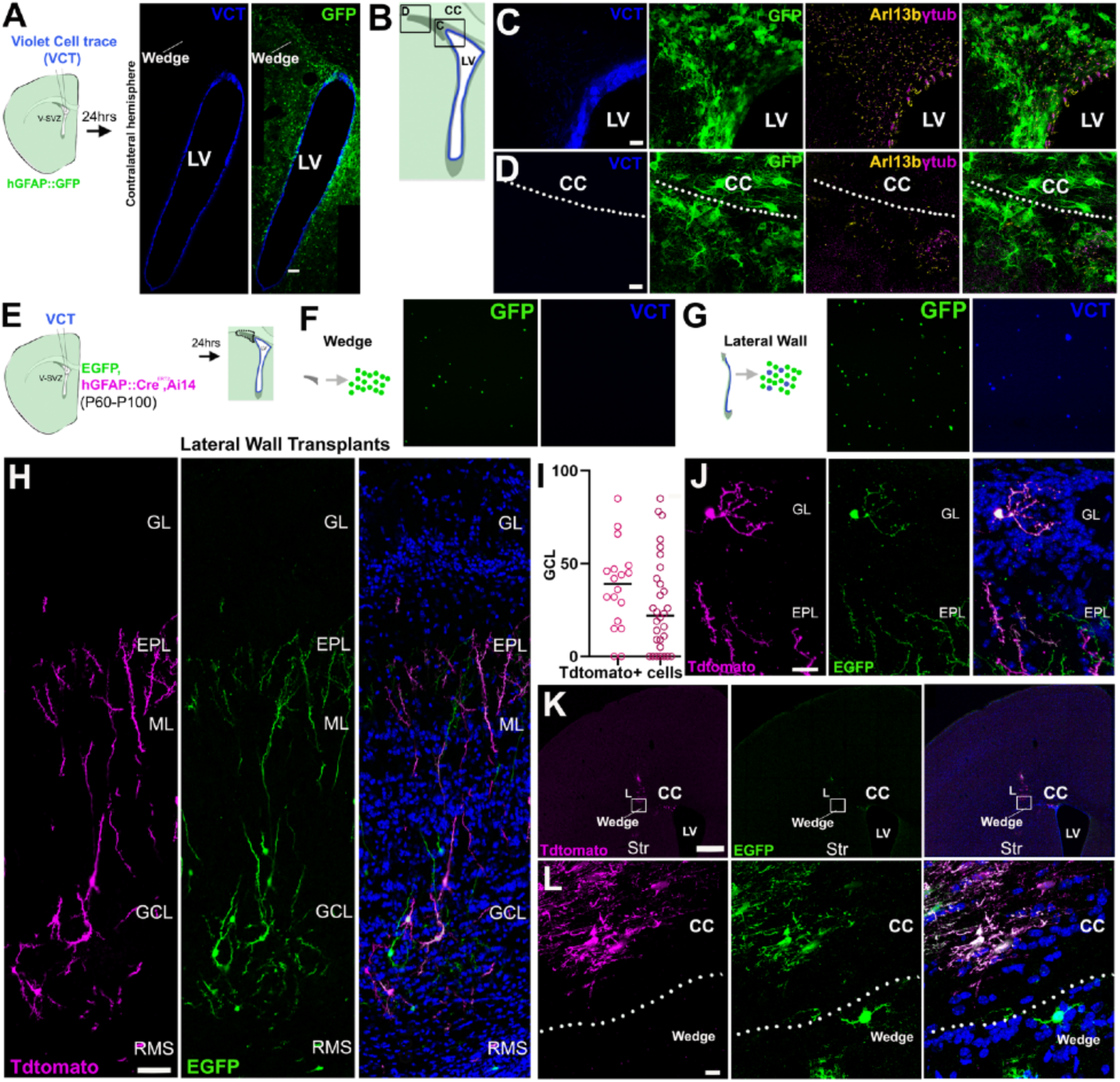
(**A-D**) Confocal images of violet cell trace (VCT) labeling apical cells in the contralateral hemisphere of a hGFAP:GFP mouse. (**C-D**) B2 cells identified by GFP expression (green) and the localization of the primary cilium (Arl13b, yellow)-basal body (γ-tubulin, magenta) were VCT^-^ in the dorsal lateral wall (C) and the wedge regions (D). (**E-G**) Epifluorescence images of isolated cells from microdissected wedge (F) and lateral wall (G) regions. Note that wedge cells are not labeled with VCT. (**H-L**) Confocal images of Lateral wall transplants 30 days after transplantation. (**H**) Confocal images of Tdtomato^+^/GFP^+^ granular cells in the OB of a transplanted mouse. (**I**) Granule cell layer (GCL) position of wedge transplant-derived OB granular cell interneurons per animal (symbols represent individual cells). 0 represents deep GCL. (**J**) Confocal images of periglomerular Tdtomato^+^/GFP^+^ cells. (**K-L**) Transplantation site showing Tdtomato^+^/GFP^+^ oligodendrocytes in the corpus callosum. GL: Granular layer, EPL: External plexiform layer, ML: Mitral layer, GCL: granular cell layer, RMS: rostral migratory stream, CC: corpus callosum, Str: Striatum, LV: lateral ventricle. Scale bars: 0.5mm (K), 50μm (B,H), 20μm (J), 15μm (C,D),10μm (L).

